# Post-translational modifications drive protein stability to control the dynamic beer brewing proteome

**DOI:** 10.1101/358796

**Authors:** Edward D. Kerr, Christopher H. Caboche, Benjamin L. Schulz

## Abstract

Mashing is a key step in beer brewing in which starch and proteins are solubilized from malted barley in a hot water extraction and digested to oligomaltose and free amino nitrogen. We used SWATH-MS to measure the abundance and site-specific modifications of proteins throughout a small-scale pale ale mash. Proteins extracted from the malt at low temperatures early in the mash decreased precipitously in abundance at higher temperatures late in the mash due to temperature/time-induced unfolding and aggregation. We validated these observations using experimental manipulation of time and temperature parameters in a micro-scale pale ale mash. Correlation analysis of temperature/time-dependent abundance showed that sequence and structure were the main features that controlled protein abundance profiles. Partial proteolysis by barley proteases was common early in the mash. The resulting proteolytically clipped proteins were particularly sensitive and were preferentially lost at high temperatures late in the mash, while intact proteins remained soluble. The beer brewing proteome is therefore driven by the interplay between protein solubilisation and proteolysis, which are in turn determined by barley variety, growth conditions, and brewing process parameters.

## Introduction

Beer brewing is a well-established industrial process comprising a series of connected bioprocesses (Fig. 1A). Harvested barley seeds are turned into malt through a process called malting, which involves controlled germination and drying (1, 2). Mashing then extracts soluble components from the malt and allows enzymes to degrade starch into small fermentable sugars and proteins into free amino nitrogen (FAN) (1). The soluble product of mashing is called wort, which is boiled with hops to sterilize, introduce bitterness, remove volatile off-flavours, and stop enzymatic activity (1, 3). Fermentation is the next stage of the brewing process, in which yeast ferments sugars to ethanol, producing beer. This beer can then be optionally filtered to create “bright beer”, or left unfiltered, further aged or not, carbonated, and packaged.

**Figure 1.**
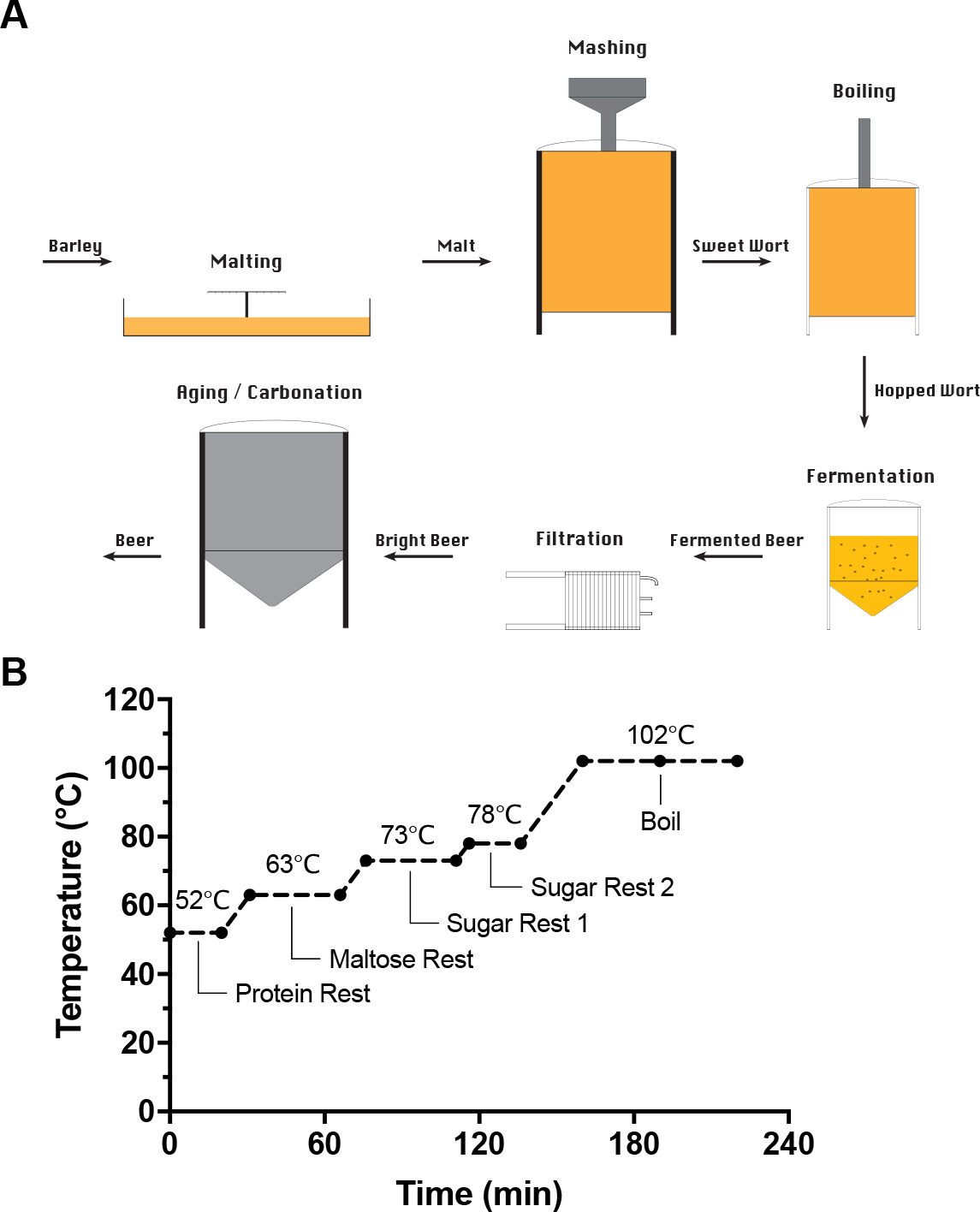
Overview of the brewing process. (**A**) Overview of the brewing process from malting to beer. (**B**) Mashing program used in this study.

While it is appreciated that the mash and boil entail complex biochemistry, the details of aspects of these processes are poorly understood. Proteomic studies have identified many proteins in wort and in beer, highlighting that some proteins can survive proteolysis in the mash and heating during malting and boiling (4–10). Studies using two-dimensional gel electrophoresis (2-DE) and mass spectrometry identified these heat-stable proteins as cysteine-rich plant defence proteins, α-amylase/trypsin inhibitors, and non-specific lipid transfer proteins (4, 5). More recent studies have shown that the wort and beer proteome is highly complex, and contains hundreds of unique proteins from barley and yeast (6–10). Complex post-translational modifications including proteolysis, oxidation, and glycation are also present on proteins in wort and beer (8, 11, 12). While these studies demonstrate the complexity of the wort and beer proteomes, precisely when during the brewing process protein modification occurs and how the wort proteome changes during the mash remain unexplored.

Proteins are extracted from malt during the mash and are subsequently denatured, aggregated, and precipitated at higher temperatures in the mash or boil (3). However, the proteome-wide extent of temperature-dependent protein aggregation during mashing has not been studied. Proteome thermal profiling has been performed of other complex systems including studying proteome-wide drug-protein interactions in mammalian cells (13–16). This approach measures the thermal stability of every detectable protein in a proteome by LC-MS quantification of the fraction of each protein that is soluble after incubation at various temperatures. The proteome thermal profiling workflow mimics the temperature ramp used in typical beer mashing and boiling, and by analogy suggests that the dynamic wort proteome is balanced by competing processes including protein solubilisation, modification, and aggregation.

In this study, we used Sequential Window Acquisition of All THeoretical Mass Spectra (SWATH-MS) to address the complexity of the dynamics of protein abundance and modification during beer brewing. We uncover an incredible complexity of protein modifications and show that an interplay of partial proteolysis and temperature-dependent protein unfolding drive the proteome of wort and beer.

## Methods

### Experimental design and statistical rationale

A small-scale mash was performed using a Braumeister (Speidel) with 23 L of water and 5 kg of milled Vienna pale ale malt (Brewer’s Choice, Brisbane) with a multi-step mash (Table S1 and Fig. 1B). Samples were taken at the start and end of each stage, and after 30 min of boil. Samples were clarified by centrifugation at 18,000 rcf at room temperature for 1 min immediately after sampling, and the supernatants stored at −20°C. Samples were taken from independent triplicate mashes using malt from the same batch. Micro-scale mashing was performed using the same multi-step mash profile as the small-scale mash with additional time extensions at each stage of the mash (Table S7) to allow differentiation between temperature- and time-dependent proteomic changes. Each replicate sample was prepared individually in a 1.5 mL protein Lobind tube (Eppendorf) with 200 mg of milled pale ale malt resuspended in 1.0 mL of water. Samples were incubated in a Multi-Therm Heat-Shake incubator (Benchmark Scientific) shaking at 1500 rpm. Triplicate samples were taken at the start, end, and extension of each stage, and after 30 min of boil. The soluble fraction was separated from the grist by centrifugation at 18,000 rcf at room temperature for 1 min immediately after sampling, transferred to a new tube, and further clarified by centrifugation at 18,000 rcf at room temperature for 1 min. Proteins from 10 μL or 100 μL of each sample from small- or micro-scale mashes, respectively, were precipitated by addition of 100 μL or 1 mL 1:1 methanol/acetone, incubation at −20°C for 16 h, and centrifugation at 18,000 rcf at room temperature for 10 min. Proteins were resuspended in 100 μL 100 mM ammonium acetate and 10 mM dithiothreitol with 0.5 μg trypsin (Proteomics grade, Sigma) and digested at 37 °C with shaking for 16 h. Trypsin was added at equal amounts to all samples to allow normalisation of relative protein abundance. SWATH-MS was implemented as described below using triplicates to reduce retention time variation and improve data quality.

### SWATH-MS

Peptides were desalted with C18 ZipTips (Millipore) and measured by LC-ESI-MS/MS using a Prominence nanoLC system (Shimadzu) and TripleTof 5600 mass spectrometer with a Nanospray III interface (SCIEX) essentially as previously described (17). Approximately 1 μg or 0.2 μg desalted peptides, estimated from ZipTip peptide binding capacity, were injected for data dependent acquisition (DDA) or data independent acquisition (DIA), respectively. Peptides were separated on a VYDAC EVEREST reversed-phase C18 HPLC column (300 Å pore size, 5 μm particle size, 150 μm i.d. × 150 mm) at a flow rate of 1 μl/min) with a linear gradient of 10-60% buffer B over 14 min, with buffer A (1% acetonitrile and 0.1% formic acid) and buffer B (80% acetonitrile and 0.1% formic acid), for a total run time of 24 min per sample. LC parameters were identical for DDA and DIA, and DDA and DIA MS parameters were set as previously described (18).

### Data analysis

Peptides and proteins were identified using ProteinPilot 5.1 (SCIEX), searching against all eukaryotic proteins in UniProtKB (downloaded 29/01/2015, 547351 total entries), with settings: sample type, identification; cysteine alkylation, none; instrument, TripleTof 5600; species, none; ID focus, biological modifications; enzyme, trypsin; search effort, thorough ID. The abundance of peptide fragments, peptides, and proteins was determined using PeakView 2.1 (SCIEX), with settings: shared peptides, allowed; peptide confidence threshold, 99%; false discovery rate, 1%; XIC extraction window, 6 min; XIC width, 75 ppm. The mass spectrometry proteomics data have been deposited to the ProteomeXchange Consortium via the PRIDE partner repository with the dataset identifiers PXD010013 and PXD013177 (19). For protein-centric analyses, protein abundances were normalised to the abundance of trypsin self-digest peptides. For analysis of peptide modifications, the abundance of each variant form of a peptide was normalised to the summed abundance for all detected forms of that peptide. For micro-scale mash analysis, PeakView output was reformatted with a python script (https://github.com/bschulzlab/reformatMS) (20), and significant differences in protein abundance were determined using MSstats (2.4) in R (21), with a significance threshold of P = 0.05.

Proteins were clustered based on correlation profiling of their abundance throughout the mash. We defined a measure of distance between pairs of proteins based on how different their abundances were at each time point. The distance between protein *a* and protein *b* was given by: 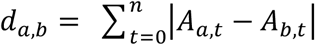, where *A*_*a,t*_ is the normalized abundance of protein *a* at time point *t*. The difference between every pair of proteins across the entire temperature profile was then calculated by summing the distance across all time points for each protein. Proteins with similar stability profiles were clustered using Cluster 3.0 (22), implementing a hierarchical, uncentered correlation, and complete linkage on both x and y. Clustered data were represented as a heat map using Plotly (1.12.2). Physicochemical properties of proteins including amino acid length, molecular weight, pI, aliphatic index, charge at pH 7, charge at pH 5, hydrophobicity, and amino acid composition was calculated using Peptides 2.2 (23). GO term enrichment was determined using GOstats (24) in R using a significance threshold of P = 0.05.

## Results

Enzymes degrade starch and proteins during the mash to produce oligomaltose sugars and FAN, while boiling the wort halts enzyme activity by denaturing proteins. We used a step mash with a protein rest (period of stationary temperature) at 52°C for 20 min, maltose rest at 63°C for 35 min, sugar rest 1 at 73°C for 35 min, sugar rest 2 at 78°C for 20 min, and finally the boil at 102°C for 60 min (Fig. 1B). This temperature ramp allowed detailed analysis of the temperature/time dependent changes in the wort proteome during the mash and boil. Liquor was sampled at the start and end of each rest and after 30 min of the boil, clarified by centrifugation, precipitated, digested by trypsin, desalted, and analysed by SWATH-MS.

We identified peptides and proteins in the samples using DDA LC-MS/MS detection and ProteinPilot identification. Protein identification, quantification, and raw mass spectrometry data are available via the PRIDE partner repository with the dataset identifier PXD010013. We identified a total of 87 unique proteins. As we searched the entire UniProt database some non-barley proteins were identified, and these likely correspond to unannotated proteins in the barley proteome. Of the proteins identified, 7 were involved in starch degradation, 21 were defence proteins, 7 were nutrient reservoirs, 12 were amylase/protease inhibitor proteins, and the majority of the remainder were house-keeping proteins.

### Global proteomic changes during mashing

After identifying peptides and proteins, we measured the abundance of each protein in each sample using SWATH-MS and PeakView, normalising protein abundance to trypsin self-digest peptides. This allowed us to determine changes in protein levels during the mash and boil, and thereby measure protein-specific extraction, solubilisation, aggregation, and precipitation of proteins during mashing. Our data showed that the abundance of most proteins increased during the maltose rest (63°C) and sugar rest 1 (73°C), and then decreased substantially during sugar rest 2 (78°C) and into the boil (102°C) (Fig. 2A). This is consistent with efficient extraction of proteins from the grain at lower temperatures early in the mash, followed by temperature/time-dependent unfolding, aggregation, and precipitation at higher temperatures later in the mash.

To understand how the relative abundance of proteins changed during the mash, and what characteristics determined the stability of a protein during the mash and boil, we visualised the relative abundance of classes of proteins that are of particular importance to barley and beer quality: the most abundant proteins, α and β-amylase, beer foam forming or associated proteins, and defence proteins. NLTP1 (non-specific lipid-transfer protein 1) was the most abundant protein as measured by MS peptide signal throughout the mash, followed by IAAD, IAAA, and IAAB (Fig. 2B). The amount of these proteins increased until the end of the first sugar rest (73°C), after which their abundance quickly declined by the end of the second sugar rest (78°C).

**Figure 2.**
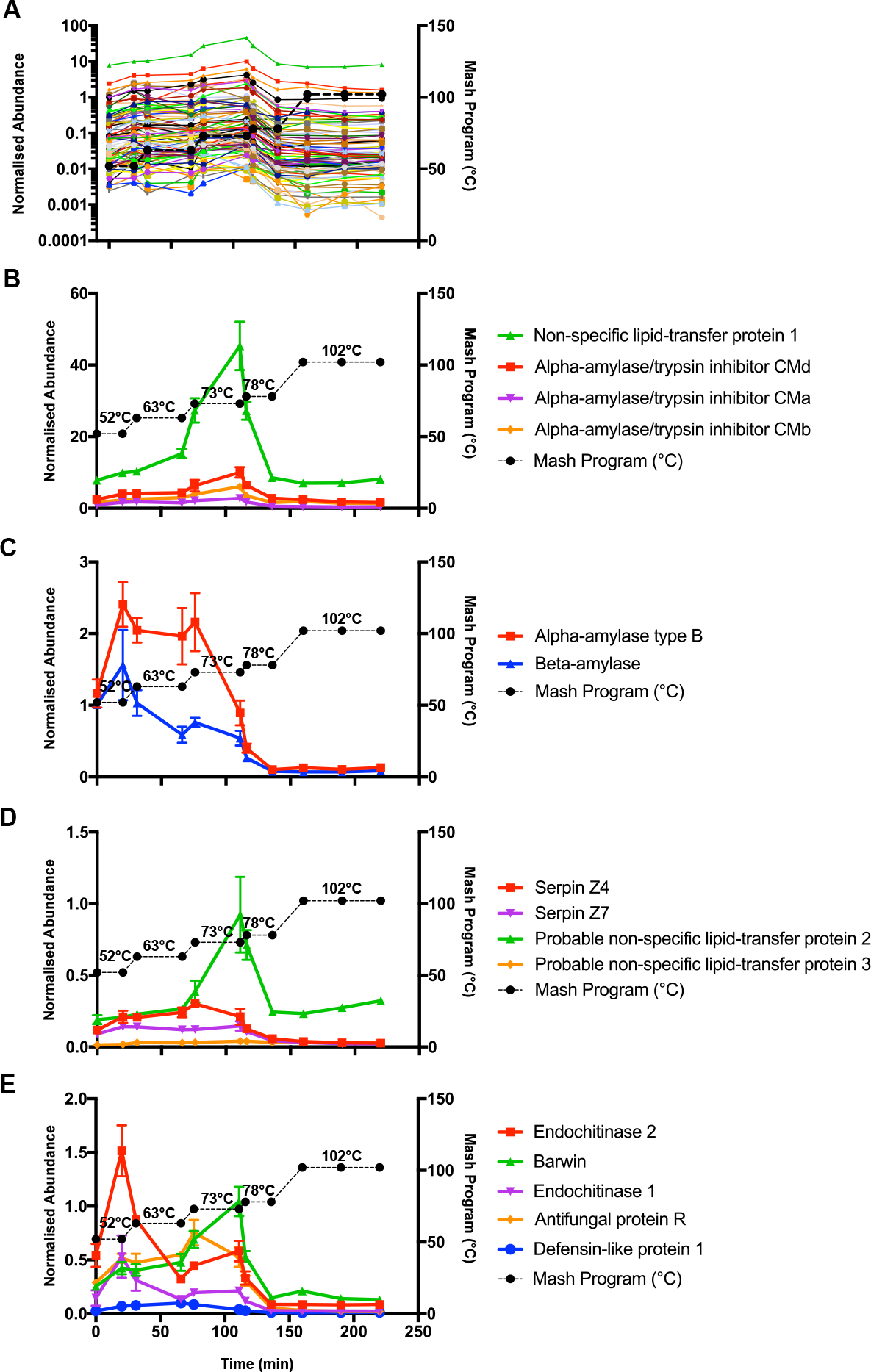
Dynamics of protein abundance during the mash and boil. (**A**) Log-scale abundance of all 87 identified and measured proteins. Error bars and protein names not shown for clarity. Each protein is represented by a different colour. (**B**) Abundance of the four most abundant proteins: α-amylase/trypsin inhibitor CMd (IAAD), non-specific lipid-transfer protein 1 (NLTP1), α-amylase/trypsin inhibitor CMa (IAAA), and α-amylase/trypsin inhibitor CMb (IAAB). (**C**) Abundance of α-amylase type B (AMY2) and β-amylase (AMYB). (**D**) Abundance of beer foam formation and stability proteins: Serpin Z4, Probable non-specific lipid-transfer protein (NLTP2), Serpin Z7, and Probable non-specific lipid-transfer protein (NLTP3). (**E**) Abundance of selected defence proteins: endochitinase 2 (CHI2), barwin (BARW), endochitinase 1 (CHI1), antifungal protein R (THHR), and defensin-like protein 1 (DEF1). A – E: Abundance (a.u; arbitrary units) of each protein normalised to trypsin. Values show mean, n=3. Error bars show SD. Mash temperature profile is shown on right Y-axis.

α- and β-amylase are critical for the hydrolysis of starch into fermentable sugars by cleaving starch at internal α(1, 4) linkages and at non-reducing ends, respectively (1, 3, 25). The activities of these enzymes *in situ* during the mash have been reported to peak at 63°C for both α and β-amylase, and decline above 70°C for α-amylase and at 65°C for β-amylase (26, 27). Our proteomic analysis showed that α- and β-amylase initially had similar levels in the mash (Fig. 2C). The abundance of α-amylase increased during the protein rest (52°C) and was stable until the first sugar rest (73°C) where it slowly declined to be essentially absent at the end of the second sugar rest (78°C). β-amylase followed a similar trajectory but at consistently lower abundance. The temporal changes in the levels of these proteins we observed here is therefore consistent with the temperature-dependent enzyme activities (26, 27), consistent with soluble enzyme abundance driving overall amylase activity in wort.

We detected several proteins which have been reported to play roles in beer foam formation and stability: non-specific lipid-transfer proteins 1 – 3 (NLTPs), Serpin Z4, and Serpin Z7 (Fig. 2B, D). Lipid-transfer proteins (LTPs) and serpin proteins are reported to be positively associated with beer foam formation and stability (28–30). LTPs all increased until the end of the maltose rest (63°C) then decreased once temperature rose above 63°C (Fig. 2B, D). Serpin proteins increased until the maltose rest (63°C), then decreased (Fig. 2D). Interestingly, a substantial fraction of NLTP1 and NLTP2 were still abundant throughout the boil and into the final beer (Fig. 2B, D).

Anti-fungal defence proteins were abundant throughout the mash (Fig. 2E), consistent with previous proteomic studies of barley and beer (4–6, 31, 32). We therefore examined how key defence proteins changed during the mash (Fig. 2E). These proteins were all efficiently extracted at low temperatures early in the mash, and then showed diverse abundance profiles throughout the mash. At the beginning of the mash, 26 kDa endochitinase 2 (CHI2) was the most abundant defence protein, but its abundance decreased at the maltose rest (63°C). 26 kDa endochitinase 1 (CHI1) followed a similar trend, but at a consistently lower abundance (Fig. 2E). While levels of CHI1 and CHI2 decreased during the first sugar rest (73°C), Barwin (BARW) and Antifungal Protein R (THHR) increased during this stage. The abundance of BARW and THHR then decreased until the end of the second sugar rest (78°C). Defensin-like protein 1 (DEF1) had a low abundance throughout the mash compared to other defence proteins but increased during the maltose (63°C) and first sugar rest (73°C). Defence proteins were generally amongst the most stable proteins, as they were present at high abundance until the first sugar rest (73°C), and were still present at moderate levels at the end of the boil.

### Proteomic changes during mashing depend on time and temperature

While protein abundances changed throughout the mash, it was not clear if these changes were dependent on the dynamics of temperature or time. To investigate these factors in more detail we performed a micro-scale mash that replicated the conditions of the small-scale (20 L) mash, but that more easily allowed manipulation of mashing conditions. We implemented a micro-scale mash in a benchtop thermomixer at millilitre scale, and then compared the proteomes throughout mashing of a standard temperature program and programs with continued incubation at each temperature stage (Fig. 3). This experimental design allowed us to discriminate between the contributions of temperature and time to changes in the mashing proteome.

We first compared the proteomic profile of NLTP1 and IAAA during the micro-scale (1.0 mL) mash (Fig. 3) and in the small-scale (20 L) mash (Fig. 2B), and found that these were qualitatively very similar. We next tested if changes in the abundance of NLTP1 and IAAA during the micro-scale mash were dependent on time, temperature, or on a combination of both factors. For NLTP1, we found that its increase during the maltose rest (63°C) was primarily time-dependent, as NLTP1 showed significantly increased abundance after further incubation at 52°C or 63°C (Fig. 3A, B). Similarly, its decrease after the maltose rest (63°C) and during sugar rest 1 (73°C) was predominantly time-dependent, as NLTP1 showed significantly decreased abundance after further incubation at 63°C or 73°C (Fig. 3A, B). Following this, the decrease in protein abundance during sugar rest 2 (73°C) was primarily temperature-dependent, as NLTP1 showed significantly decreased protein abundance after incubation at 78°C compared with further incubation at 73°C (Fig. 3A, B).

**Figure 3.**
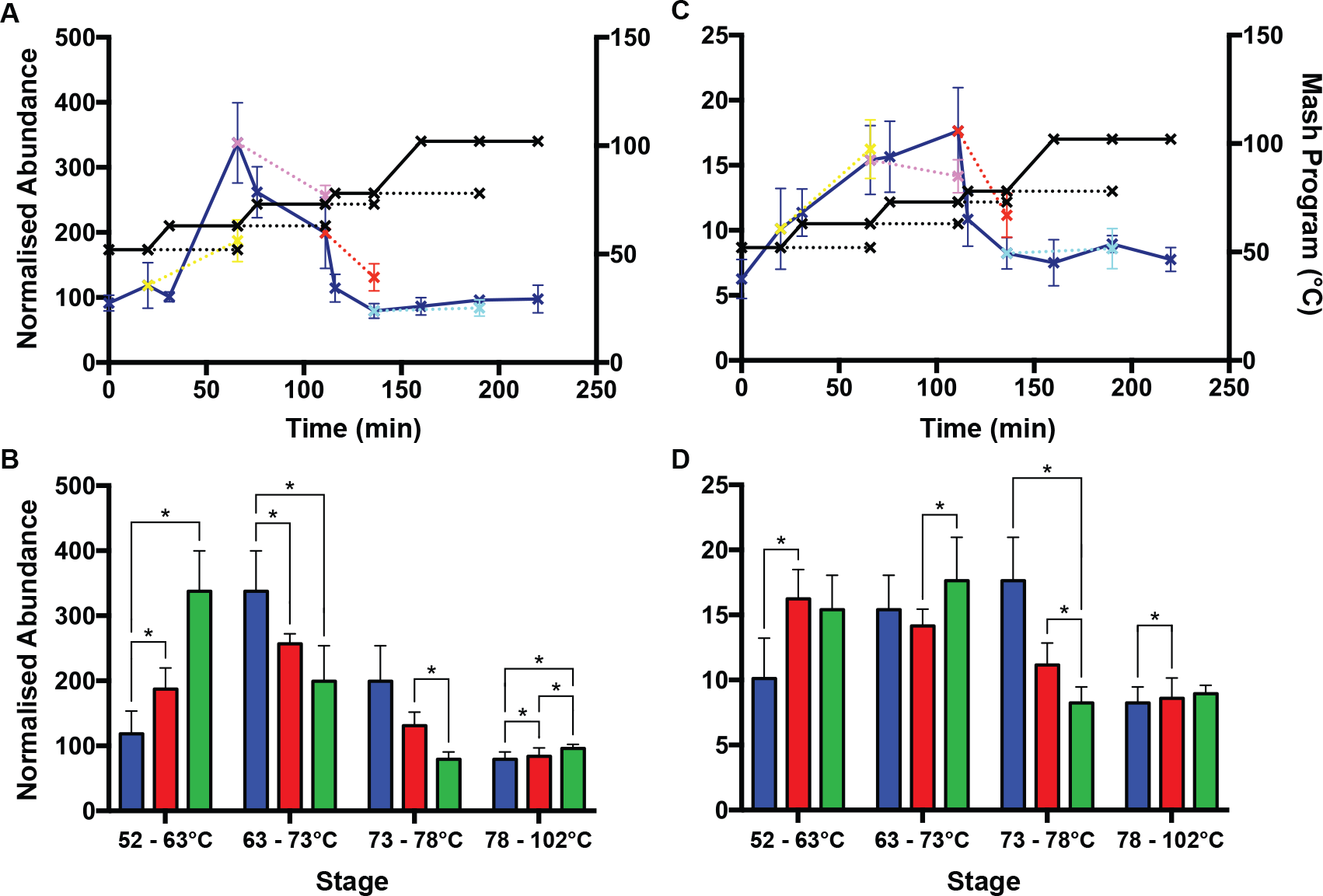
Changes in protein abundance dependent on time or temperature. Normalized abundance of **(A)** non-specific lipid-transfer protein 1 (NLTP1) and **(C)** α-amylase/trypsin inhibitor CMa (IAAA). Standard mash program, solid line, dark blue; rest stage temperature extensions, dotted lines, yellow (52°C), purple (63°C), red (73°C), and cyan (78°C). Mash temperature profile is shown on the right Y-axis. Mash program is shown in black solid and dotted lines; crosses indicate when samples were taken. Normalized abundance of **(B)** NLTP1 and **(D)** IAAA between sequential stages. First stage (blue), time extension of first stage (red), and next temperature stage (green). *, P < 0.05. Values show mean, n=3. Error bars show SEM.

For IAAA, we found that the increase in protein abundance from the protein rest (52°C) to the maltose rest (63°C) was primarily time-dependent, as IAAA showed significantly increased abundance after further incubation at 52°C (Fig. 3C, D). The further increase in protein levels during sugar rest 1 (73°C) was predominantly temperature-dependent, as IAAA showed significant increase in protein abundance after an increase in temperature to 73°C but not with further incubation at 63°C (Fig. 3C, D). Its decrease during sugar rest 2 (78°C) was predominantly temperature-dependent, as IAAA showed significantly decreased protein abundance after incubation at 78°C, and not with further incubation at 73°C (Fig. 3C, D). In summary, our micro-mashing experiment revealed that both temperature and incubation time contributed to proteome-wide changes during the mash process.

### Correlation profiling of protein abundance throughout the mash

To further understand the dynamics of protein abundance throughout the mash we performed protein correlation profiling. We performed a pairwise comparison between every protein at every period of the small-scale mash and then calculated a distance matrix between each pair of proteins. This distance matrix allowed the comparison of abundance profiles between proteins. We then performed a cluster alignment to group proteins with similar profiles (Fig. 4A). This analysis identified four clusters of proteins that were primarily distinguished by the temperature/time at which the proteins reached maximum abundance (Fig. 4A, B). The presence of four distinct clusters of temperature/time-dependent abundance suggested that there were sequence, physical, or biochemical features shared between proteins in each cluster. To discover these features, we calculated a variety of theoretical physicochemical properties for each protein. Proteins in cluster three were significantly larger than proteins in clusters one and two, as measured by amino acid length or molecular weight (Fig. 4C), and were enriched in aromatic amino acids (Fig. 4D). Cluster three also had the lowest temperature of maximum protein abundance of the four clusters (Fig. 4B), consistent with larger and more hydrophobic proteins being more susceptible to temperature/time-dependent unfolding and aggregation (8). Cluster one had the highest temperature of maximum abundance (Fig. 4B) and was enriched in cysteines (Fig. 4C), consistent with abundant disulfide bonds increasing protein stability.

**Figure 4.**
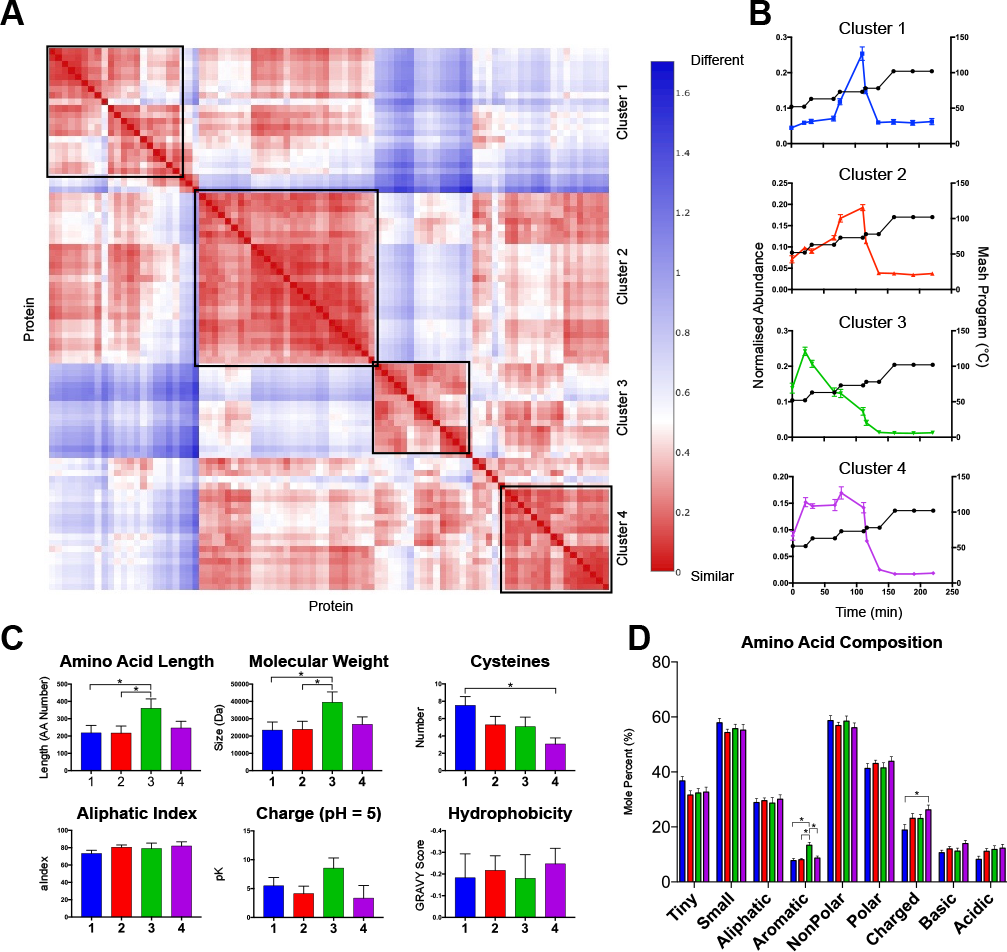
Correlation profiling of protein abundance throughout the mash. **(A)** Proteins clustered using Cluster 3.0, and heat mapped. Colour represents difference in abundance profiles of each pair of proteins: red, similar; blue, different. **(B)** cluster 1 – 4. **(C)** protein length, molecular weight, number of cysteines, aliphatic index, protein net charge at pH = 5, and hydrophobicity. Values show mean of proteins within a cluster, error bars show SEM. **(D)** Amino acid composition of proteins in each cluster. * indicate significant (p < 0.05) differences between groups.

The temperature/time-dependent abundance profiles of homologous proteins tended to be similar (Fig. 2), and each cluster of proteins with correlated abundance profiles had a unique physicochemical profile (Fig. 4). Together, this suggested that each cluster (Fig. 4A, B) consisted of homologous or related proteins. We tested this using GO term enrichment analysis on the four clusters to identify terms and pathways associated with proteins belonging to each cluster. Each cluster was significantly enriched in at least one GO term (Table S2). Cluster one was associated with lipid binding, due to enrichment of Lipid Transfer Proteins homologous to the abundant NLTP1. Cluster two was enriched in peptidase inhibition, associated with Chloroform/Methanol soluble proteins homologous to IAAA. Cluster three was enriched with hydrolase activity from amylases and chitinases. Lastly, cluster four was enriched in the generic GO term catalytic activity, from several functionally diverse cytoplasmic proteins. In summary, this showed that proteins in clusters one to three were enriched in homologous proteins likely to share sequence similarities, consistent with their correlated temperature/time-dependent abundance profiles. Together, these analyses showed that protein sequence and structural characteristics were the primary factors that controlled temperature/time dependent protein abundance profiles during the mash.

### Protein modification drives stability

While all proteins decreased in abundance at higher temperatures late in the mash, there were many proteins for which a substantial fraction survived the boil (e.g. NLTP1, NLTP2, BARW, CHI1; Fig. 2B, D, E). These proteins would potentially remain in the final beer, and have indeed been detected in beer (6–8, 33). This clear distinction in sensitivity between pools of the same protein suggested that covalent modifications may be driving diversity of protein structure and behaviour. To understand what allowed this small fraction of proteins to survive the boil we investigated proteome-wide changes in the abundance of site-specific modifications during the mash and boil.

Protein modifications including oxidation, glycation, and proteolytic cleavage are common in brewing, and impact beer product quality (8, 12, 34–37). We identified many sites with these modifications in proteins in wort. The thermal stability of proteins and their ability to survive the boil intact could be altered by these modifications. To test this, we investigated the dynamics of site-specific proteolysis and chemical modification in proteins during the mash and boil.

The barley proteases responsible for proteolysis during mashing are active at lower temperatures, where they cleave proteins during the protein rest (52°C) and add FAN to the wort (1, 3, 25, 35, 38, 39). As well as creating FAN, proteolysis influences protein stability. To monitor proteolysis during the mash we first identified non- or semi-tryptic peptides in the wort throughout the mash and boil. Not unexpectedly, proteolysis was a very common modification of proteins identified throughout the mash, with 59% of peptides detected as non- or semi-tryptic peptides. Inspection of homology models of protein structures showed that all sites of proteolysis we detected could be mapped to the surface of the substrate protein (Fig. 5, 6, Fig. S1 – S3). This is consistent with proteolytic enzymes acting on otherwise intact and folded proteins in the grain or wort, and proteolysis being limited to a few events for each individual protein.

Accurate determination of the effect of proteolysis on protein stability required high peptide coverage, and so we examined proteolysis in proteins that had many peptides identified: BARW, NLTP1, α-amylase/trypsin inhibitor CMa (IAAA), α-amylase/trypsin inhibitor CMb (IAAB), α-amylase/trypsin inhibitor CMd (IAAD), and α-amylase (AMY2). We grouped semi-tryptic derivatives of the same full tryptic peptide, and normalised their abundances to the sum of their total intensities at each time point during the mash and boil. For all peptides analysed, semi-tryptic derivatives increased and were dominant during the protein rest (52°C) and the beginning of the maltose rest (63°C) (Fig. 5, 6, Fig. S1–S3), consistent with abundant protease activity at these temperatures (1, 3, 25, 35, 38, 39). However, when the mash temperature rose above 63°C, these semi-tryptic peptides decreased in abundance relative to the full tryptic form (Fig. 5, 6, and Fig. S1 – S3). Typically, this occurred at the same temperature/time at which the overall amount of that protein decreased in the wort (Fig. 2A). We interpreted this effect as corresponding to proteolytically clipped forms of proteins unfolding and aggregating at temperatures at which the intact form of the protein remained stable. For example, most semi-tryptic derivatives of K-M_36_KPCLTYVQGGPGPSGECCNGVR_58_-D and K-C_99_NVNVPYTISPDIDCSR_115_-I from NLTP1 were more abundant at 52°C than the full tryptic form, and above 73°C these cleaved forms drastically decreased (Fig. 5C, D, E). This pattern was typical of most sites of proteolysis that could be confidently examined (Fig. 5, 6, and Fig. S1 – S3).

**Figure 5.**
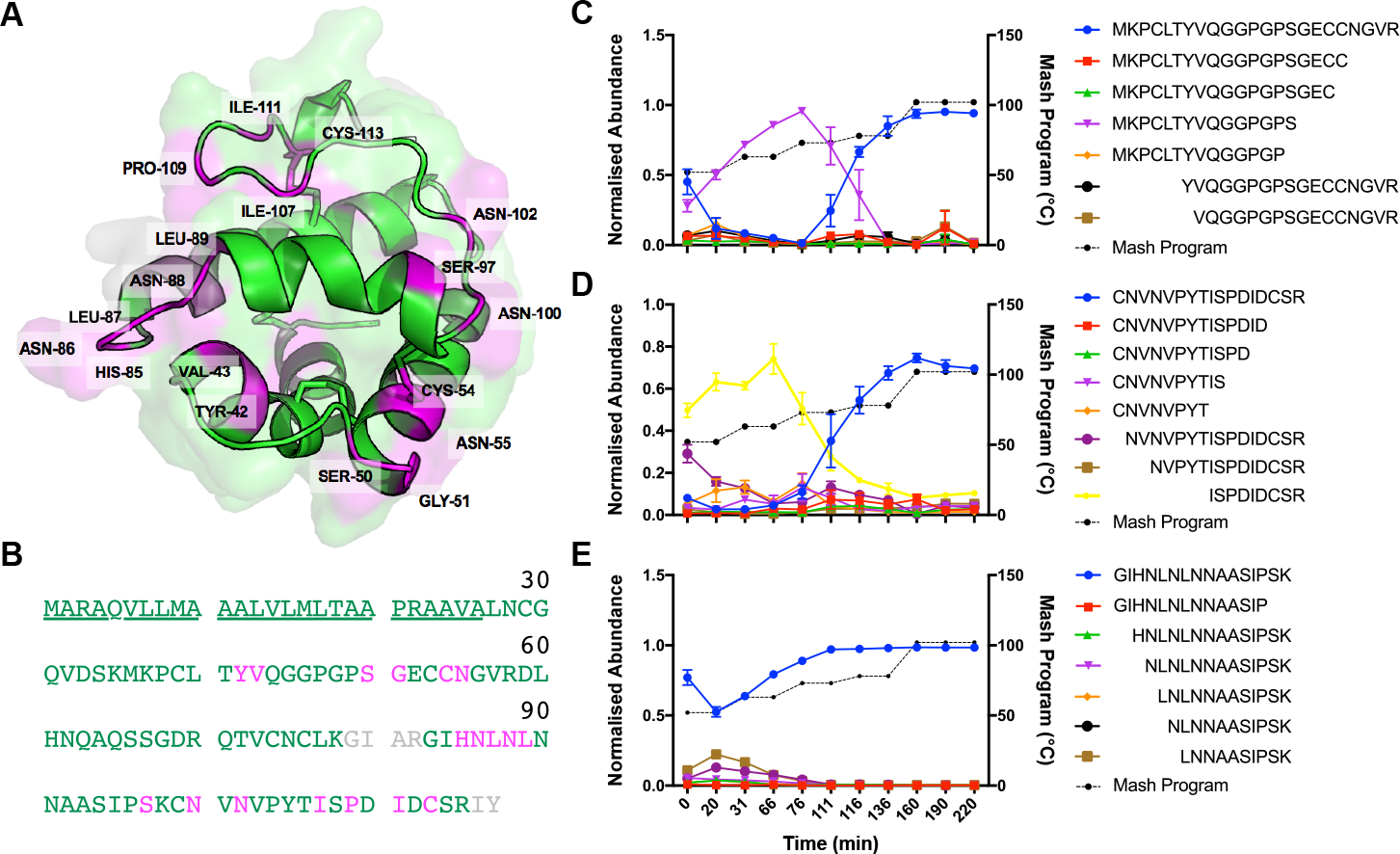
Proteolysis during the mash lowers the stability of NLTP1. **(A)** Cartoon of the X-ray crystal structure of NTLP1 (1MID). Green, peptides identified by MS; grey, unidentified peptides; magenta and labelled, amino acid after a proteolytic cleavage event. **(B)** NLTP1 protein sequence with the same colour scheme as (A). Underlined, not present in structure. Normalised abundance (a.u; arbitrary units) of full- and semi-tryptic peptides corresponding to **(C)** K-M_36_KPCLTYVQGGPGPSGECCNGVR_58_-D, **(D)** K-C_99_NVNVPYTISPDIDCSR_115_-I, and **(E)** R-G_83_IHNLNLNNAASIPSK_98_-C. Values show mean, n=3. Error bars show SEM. Mash temperature profile is shown on the right Y-axis. Mash program is shown in black dotted lines; points indicate when samples were taken.

Interestingly, a few select proteolytic cleavage events did not affect temperature/time-dependent protein abundance (Fig. 6D, S1D, and S2E). In IAAA, two semi-tryptic derivatives of K-V_122_LVTPGQCNVLTVHNAPYCLGLDI_145 (C-terminus)_ (with cleavage between H_135_-N_136_ or between D_144_-I_145_) were stable at higher temperatures and were even more abundant than the full tryptic peptide (Fig. 6). We inspected a model of the structure of IAAA to provide insight into why proteolysis at these sites might be tolerated. Disulfide bonds between C_140_ and C_87_, and between C_129_ and C_75_, likely lock the local protein fold in place to provide stability even after cleavage at H_135_-N_136_. In contrast, the C-terminal 2 amino acid residues in the model of IAAA are not defined, suggesting this region is structurally flexible. Proteolysis at D_144_-I_145_ resulting in loss of a single amino acid residue in such a flexible region is also not likely to impact protein stability. Similar effects were also observed for the N-terminal flexible extensions of IAAB and IAAD (Fig. S1D, S2E).

**Figure 6.**
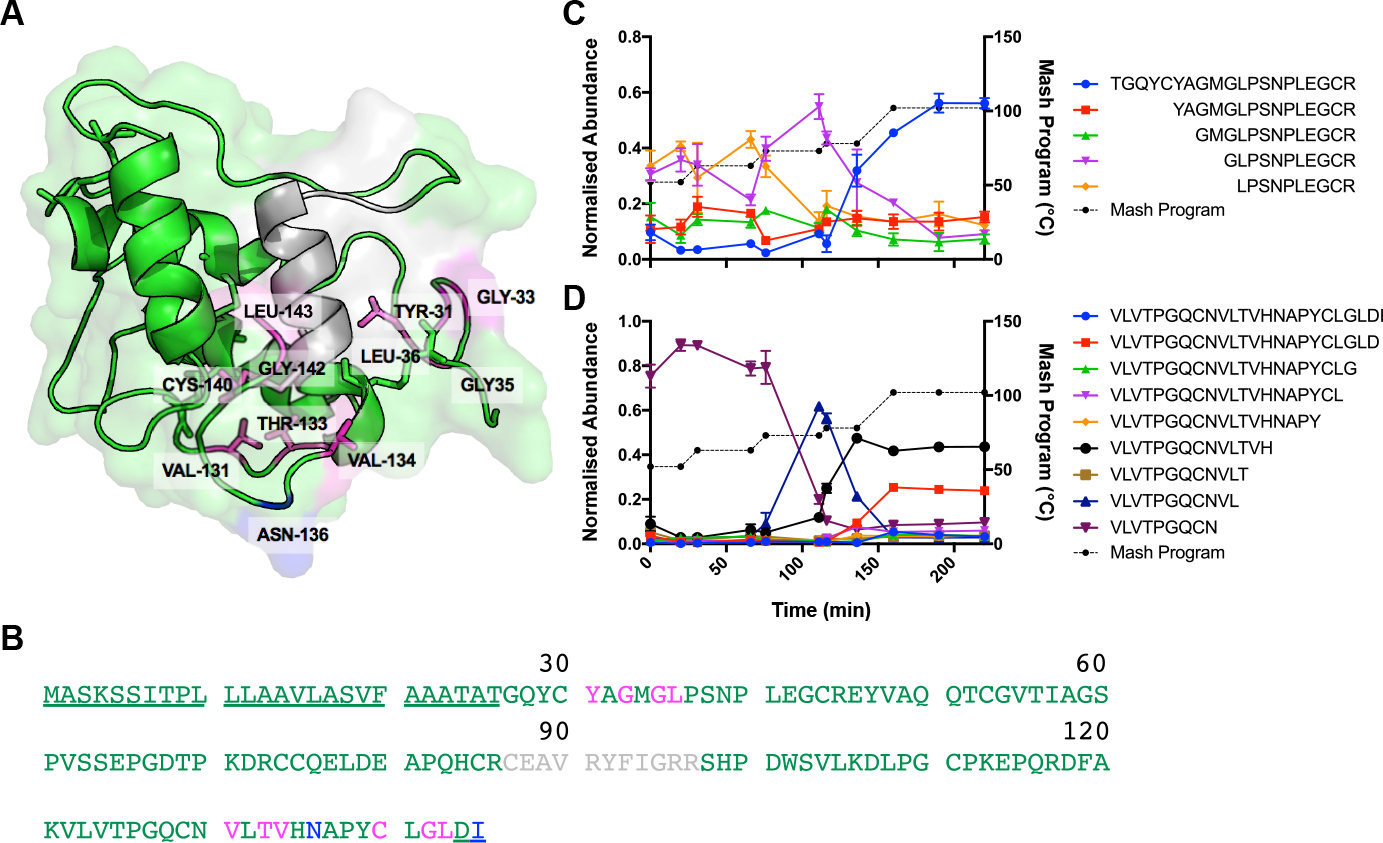
Select sites of proteolysis do not affect IAAA stability. **(A)** Cartoon of the X-ray crystal structure of IAAA, homology modelled on 1B1U (40). Green, peptides identified by MS; grey, unidentified peptides; magenta and labelled, amino acid after a proteolytic cleavage event; blue and labelled, amino acid after a proteolytic cleavage event that does not lower stability. **(B)** IAAA protein sequence with the same colour scheme as (A). Underlined, not present in structure. Normalised abundance (a.u; arbitrary units) of full- and semi-tryptic peptides corresponding to **(C)** T_26_GQYCYAGMGLPSNPLEGCR_45_-E and **(D)** K-V_122_LVTPGQCNVLTVHNAPYCLGLDI_145_. Values show mean, n=3. Error bars show SEM. Mash temperature profile is shown on the right Y-axis. Mash program is shown in black dotted lines; points indicate when samples were taken.

In summary, our data showed that extensive proteolysis during the early stages of the mash lowers the unfolding temperature of diverse proteins and explains why a large fraction of each protein unfolds, aggregates, and is lost from solution at higher temperatures. Those proteins which escape proteolysis, or which are clipped at specific sites that do not affect thermal stability, remain present even after the boil, explaining the sustained presence of a fraction of each protein through the boil and potentially into beer.

In addition to proteolysis, we also identified modification of diverse proteins by oxidation, glycation, and deamidation. Oxidation has been reported on selected proteins in beer and found to decrease as temperature increased (37). Glycation is driven by the Maillard reaction, where reducing sugars modify primarily lysine residues, catalysed by high temperature (12, 34, 41). The mash and boil therefore provide a perfect environment for glycation to take place. Although we could identify diverse sites of these modifications, data quality was insufficient to quantitatively monitor their abundance throughout the mash.

## Discussion

We used SWATH-MS to investigate the changes in the wort proteome throughout the mash and boil stages of beer brewing. Our results showed that the mash proteome was highly dynamic, with proteins increasing in abundance as they were extracted from the malt early in the mash, and then rapidly decreasing in abundance at higher temperatures later in the mash due to thermal denaturation, aggregation, and precipitation (Fig. 2). Validation with a micro-scale mash revealed that changes in protein abundance were dependent on a complex mixture of time, temperature, or a combination of both factors (Fig. 3 and Fig. S29).

Correlation profiling of protein abundance throughout the mash identified four clusters of proteins with distinct temperature/time-dependent abundance profiles and distinct physiochemical properties (Fig. 4). Each of these clusters were defined by the presence of multiple protein homologs with similar sequence and structure. Our data showed that these sequence characteristics were clearly critical in determining protein levels through the mash and into beer. However, it is likely that other process parameters such as altered germination extent during malting, grist particle size from differential grinding, or extraction efficiency through varied grist:liquor ratios would also have substantial effects on protein abundance profiles

The proteome of the soluble wort throughout the mash and boil was extremely complex, with many proteins undergoing diverse modifications. Many of these modifications functionally affected protein stability and thus abundance within the wort over the temperature profile of the mash and boil. We identified abundant proteolysis during the early stages of the mash, and correlated proteolysis with low thermal stability and specific loss of clipped proteins from the wort at higher temperatures. Extensive proteolysis and subsequent thermal destabilisation explained why a large fraction of each protein was denatured and removed from the soluble wort at higher temperatures. A small fraction of each protein generally remained soluble after the boil, due to a lack of proteolysis or only being clipped at specific sites that did not perturb thermal stability.

The proteome of the wort during the mash and boil is highly complex. Protein abundance during the mash is reliant on chemical and biophysical processes including solubilisation, proteolysis, oxidation, glycation, unfolding, and aggregation. Protein structure and stability determines whether a protein survives intact throughout the mash and boil to be present in the final soluble wort, and protein structure and stability is in turn directly affected by modifications, primarily proteolysis. Our data showed that extensive proteolytic cleavage explained why a large percentage of proteins were lost from the soluble wort during the mash and boil. Generally, proteolysis caused structural instability at high temperatures later in the mash, leading to unfolding, aggregation, and precipitation (Fig. 5, 6, Fig. S1 – S3). However, a few specific proteolytic events did not affect protein stability. These were present in unstructured terminal regions of proteins, or close to networks of disulphide bonds that would be expected to lock the local protein fold. These specific events left the proteins with sufficient stability to survive the boil intact. Interestingly, our data showed no correlation between the extent or ease of proteolysis and the subsequent impact of specific proteolytic events on protein stability. This suggests that the extent of proteolysis and the likelihood of subsequent protein unfolding are largely independent. Proteolysis is likely driven by substrate accessibility and the presence of specific recognition motifs, while unfolding of proteolytically clipped proteins is less predictable and highly dependent on local protein context.

A critical concern of the brewing industry is efficient digestion of starch to oligomaltose, which requires maintaining high activity of both α- and β-amylase throughout the mash. α-Amylase cleaves internal α(1-4) glucosidic bonds in starch, while β-amylase releases maltose disaccharides from the non-reducing ends of starch polymers or maltooligosaccharides (1, 3, 25). Maintaining high α- and β-amylase activity is made difficult by the low unfolding temperature of β-amylase relative to α-amylase (3, 26, 27). It is generally accepted that a mash temperature of ~65°C is appropriate to keep both enzymes stable while allowing efficient conversion of starch to oligomaltose, but high enough to allow adequate gelatinization of starch (1, 3). Our results confirmed that a mash temperature of ~65°C is appropriate for these aims, as both α- and β-amylase were stable at the 63°C maltose rest (Fig. 2C). Another key concern in industrial brewing is production of sufficient FAN to ensure robust yeast growth during fermentation. Our data demonstrated that proteolysis was extensive at lower temperatures (Fig. 5, 6, Fig. S1 – S3). Use of a mash profile with a low temperature stage such as a protein rest at 53°C is therefore required to achieve high FAN. However, our data also clearly demonstrated the challenges in balancing adequate FAN production with efficient starch degradation, as increased proteolysis would result in more proteolytically clipped α- and β-amylase with reduced enzymatic activity and lower thermostability. The dynamic brewing proteome and metabolome are therefore highly dependent on the abundance and structure of the starch, enzymes, and other proteins in the malt; the precise mash profile used; and the complex interplay between protein post-translational modification, activity, and stability.

Proteomics can measure aspects of the complex protein biochemistry of beer brewing inaccessible to other methods. Similar analyses could therefore be used to optimize mashing profiles for specific barley varieties, brewery processes, or beer styles based on the abundance of amylases, proteases, and other proteins. Our data also suggest that proteomics could be used for quality control of barley and malt, and to monitor process efficiency and product quality in other food bioprocessing industries.

## Acknowledgements

We gratefully acknowledge the assistance of Dr Amanda Nouwens and Mr Peter Josh at The University of Queensland School of Chemistry and Molecular Biosciences Mass Spectrometry Facility. BLS was funded by an NHMRC Career Development Fellowship. EDK was funded by an Advance Queensland PhD Scholarship.

