## Supplementary Material for "Post-translational modifications drive protein stability to control the dynamic beer brewing proteome"

**A**

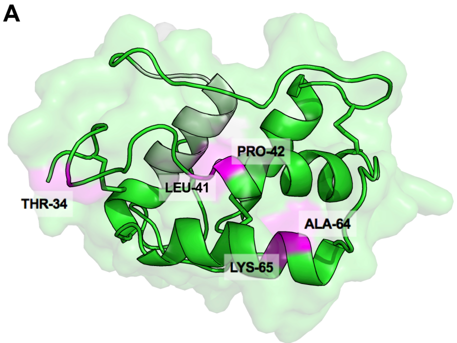


**C** **D**

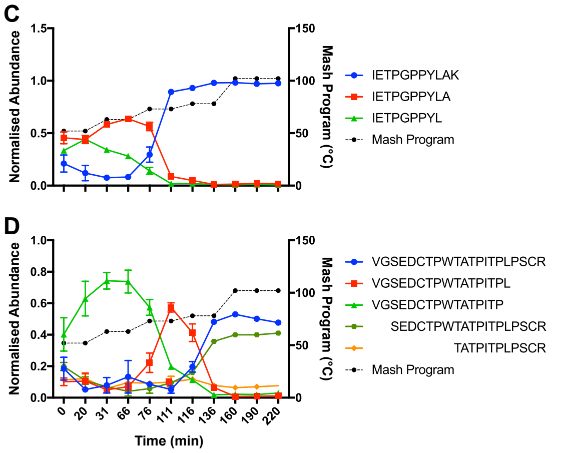


**B**

```
                                     30
MASKSSCDLL LAAVLVSIFA AVAAVGSEDC
                                     60
TPWTATPITP LPSCRDYVEQ QACRIETPGP
                                     90
PYLAKQQCCG ELANIPQQCR CQALRFFMGR
                                    120
KSRPDQSGLM ELPGCPREVQ MDFVRILVTP

GFCNLTTVHN TPYCLAMDEW QWNRQFCSS
```

**Supplementary Figure 1. Select sites of proteolysis do not affect IAAB stability. (A)** Cartoon of the X-ray crystal structure of IAAB, homology modelled on 1B1U (38). Green, peptides identified by MS; grey, unidentified peptides; magenta and labelled, amino acid after a proteolytic cleavage event. **(B)** IAAB protein sequence with the same colour scheme as (A). Underlined, not present in structure; blue, amino acid after a proteolytic cleavage event that does not lower stability. Normalised abundance (a.u; arbitrary units) of full- and semi-tryptic peptides corresponding to **(C)** R-$I_{55}$ETPGPPYLA$K_{65}$-Q and **(D)** V-$_{25}$GSEDCTPWTATPITPLPSC$R_{45}$-D. Values show mean, n=3. Error bars show SD. Mash temperature profile is shown on the right Y-axis. Mash program is shown in black dotted lines; points indicate when samples were taken.

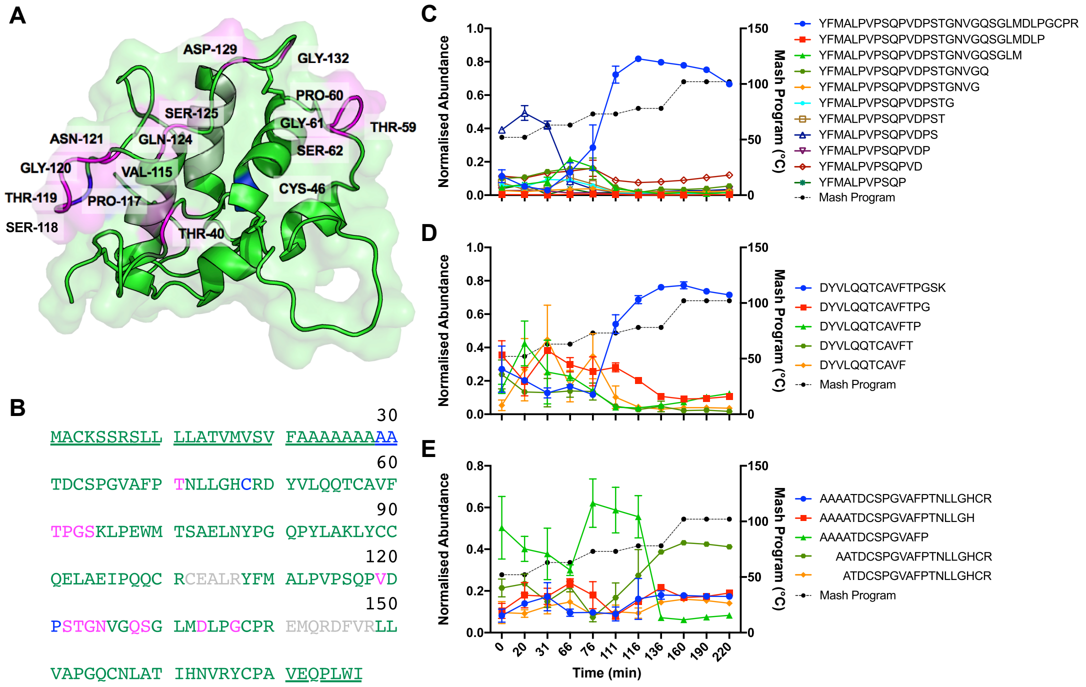


**Supplementary Figure 2. Select sites of proteolysis do not affect IAAD stability. (A)** Cartoon of the X-ray crystal structure of IAAD, homology modelled on 1B1U (38). Green, peptides identified by MS; grey, unidentified peptides; magenta and labelled, amino acid after a proteolytic cleavage event. **(B)** IAAD protein sequence with the same colour scheme as (A). Underlined, not present in structure; blue, amino acid after a proteolytic cleavage event that does not lower stability. Normalised abundance (a.u; arbitrary units) of full- and semi-tryptic peptides corresponding to **(C)** R-$Y_{104}$FMALPVPSQPVDPSTGNVGQSGLMDLPGCP$R_{135}$-E, **(D)** $D_{48}$YVLQQTCAVFTPGS$K_{63}$-L, and **(E)** $A_{25}$AAATDCSPGVAFPTNLLGHC$R_{47}$-D. Values show mean, n=3. Error bars show SD. Mash temperature profile is shown on the right Y-axis. Mash program is shown in black dotted lines; points indicate when samples were taken.

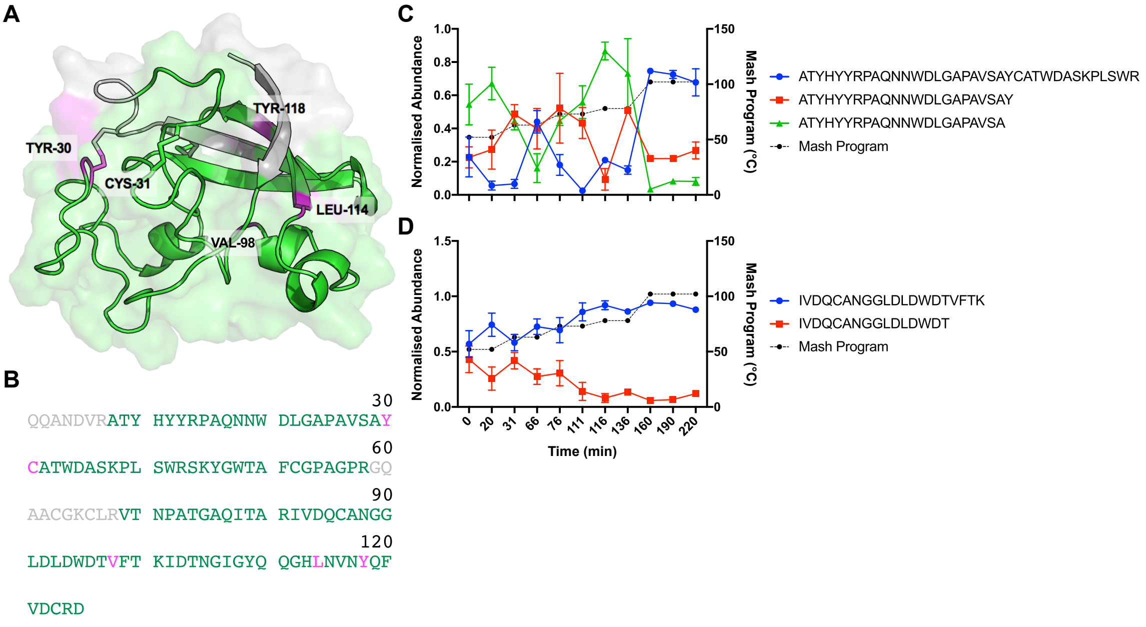


**Supplementary Figure 3. Proteolysis during the mash lowers the stability of BARW. (A)** Cartoon of the X-ray crystal structure of BARW (1BW3) (40). Green, peptides identified by MS; grey, unidentified peptides; magenta and labelled, amino acid after a proteolytic cleavage event. **(B)** BARW protein sequence with the same colour scheme as (A). Underlined, not present in structure. Normalised abundance (a.u; arbitrary units) of full- and semi-tryptic peptides corresponding to **(C)** R-$A_8$TYHYYRPAQNNWDLGAPAVSAYCATWDASKPLSWR$_{43}$-S and **(D)** R-$I_{82}$VDQCANGGLDLDWDTVFTK$_{101}$-I. Values show mean, n=3. Error bars show SD. Mash temperature profile is shown on the right Y-axis. Mash program is shown in black dotted lines; points indicate when samples were taken.

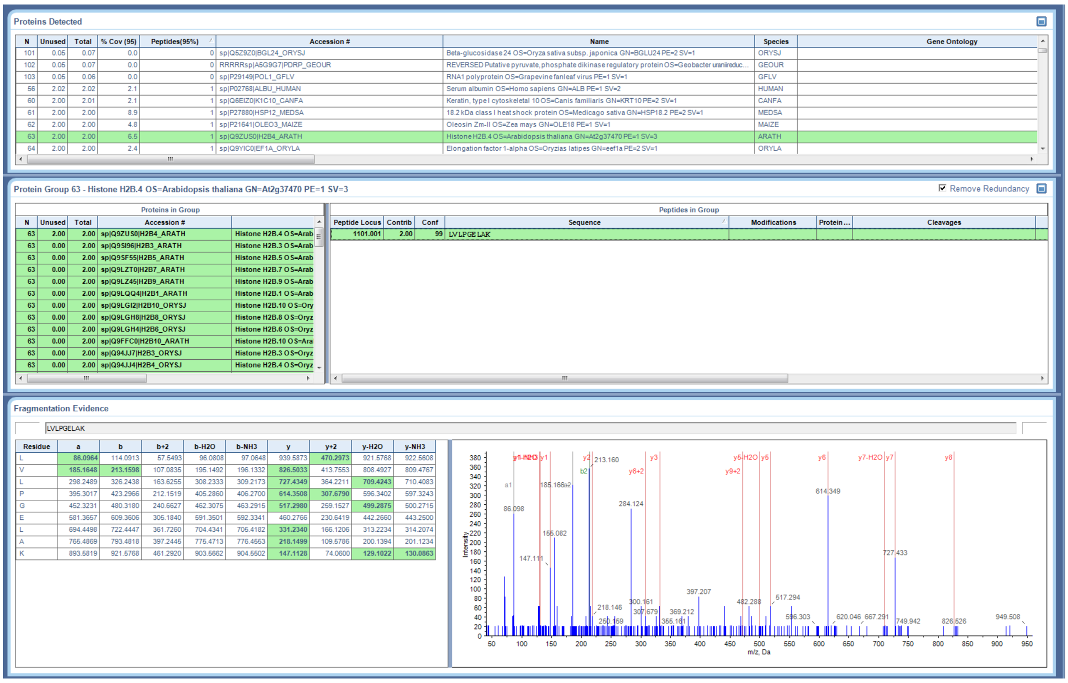


Supplementary Figure S4. MS/MS of LVLPGELAK identifying H2B4_ARATH.

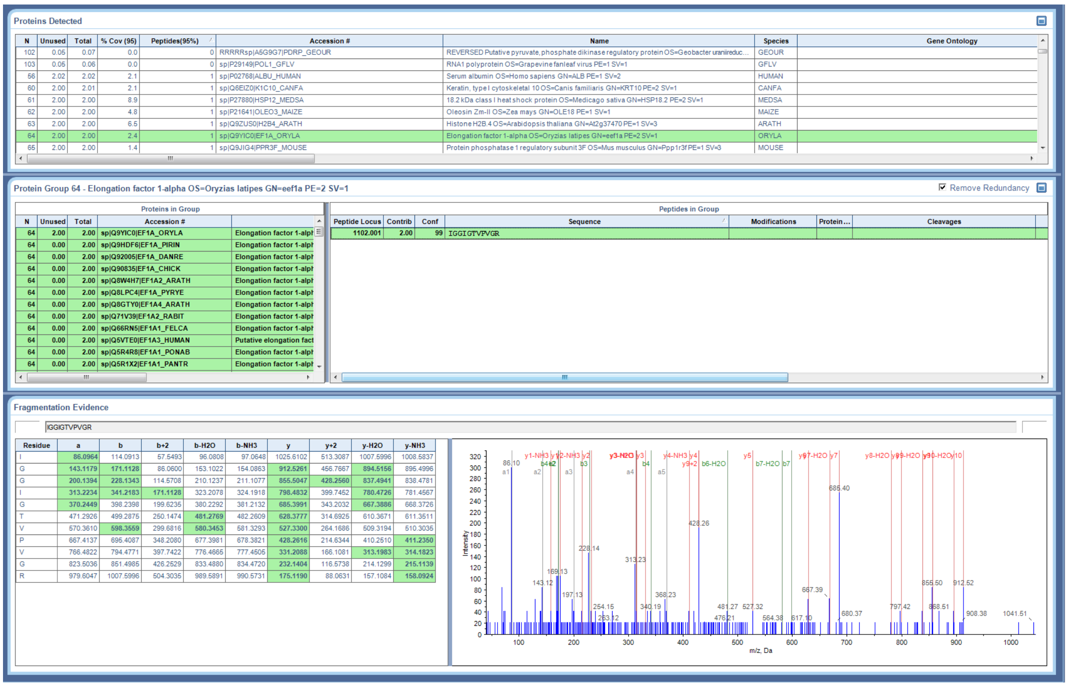


Supplementary Figure S5. MS/MS of IGGIGTVPVGR identifying ERIA_ORYLA.

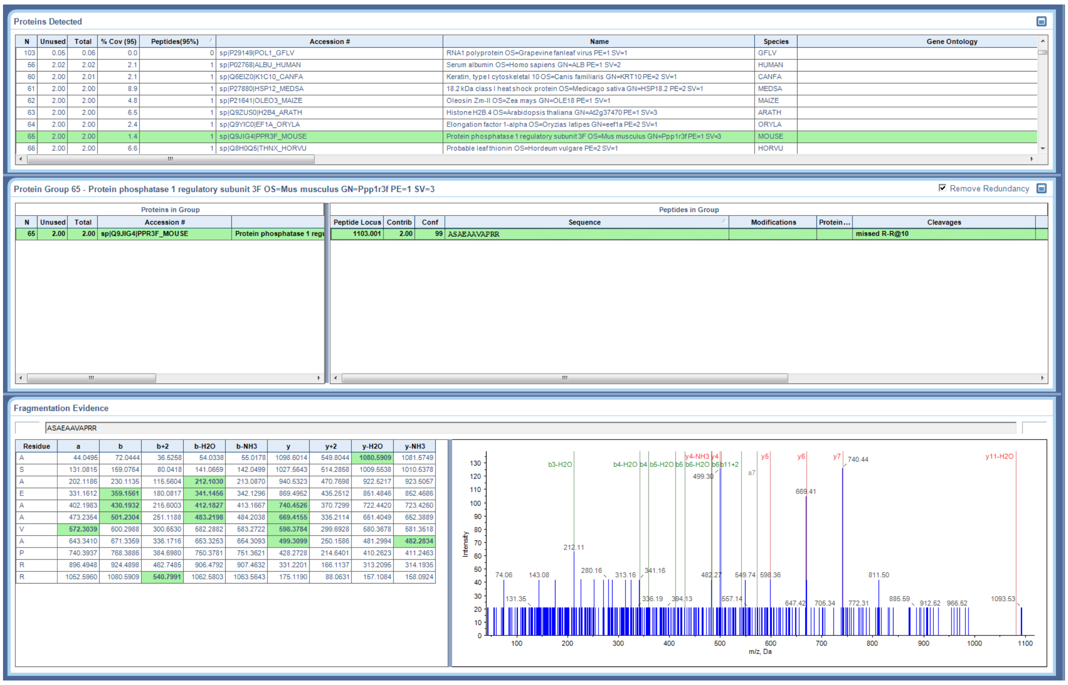


**Supplementary Figure S6. MS/MS of ASAEAAVAPRR identifying PPR3F_MOUSE.**

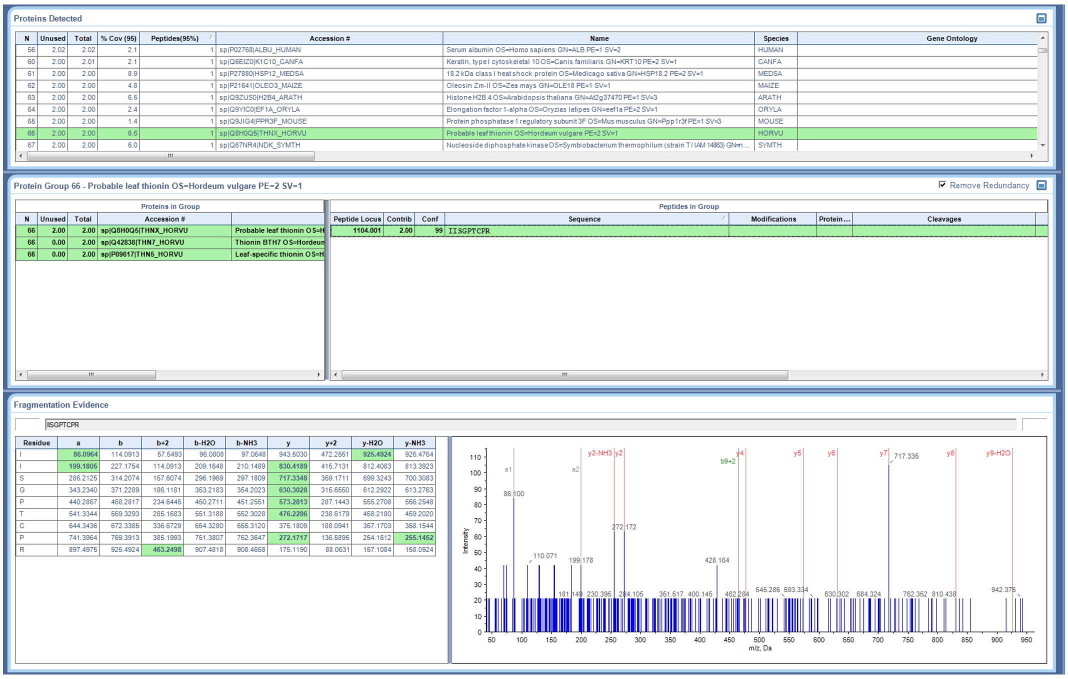


**Supplementary Figure S7. MS/MS of IISGTCPR identifying THNX_HORVU.**

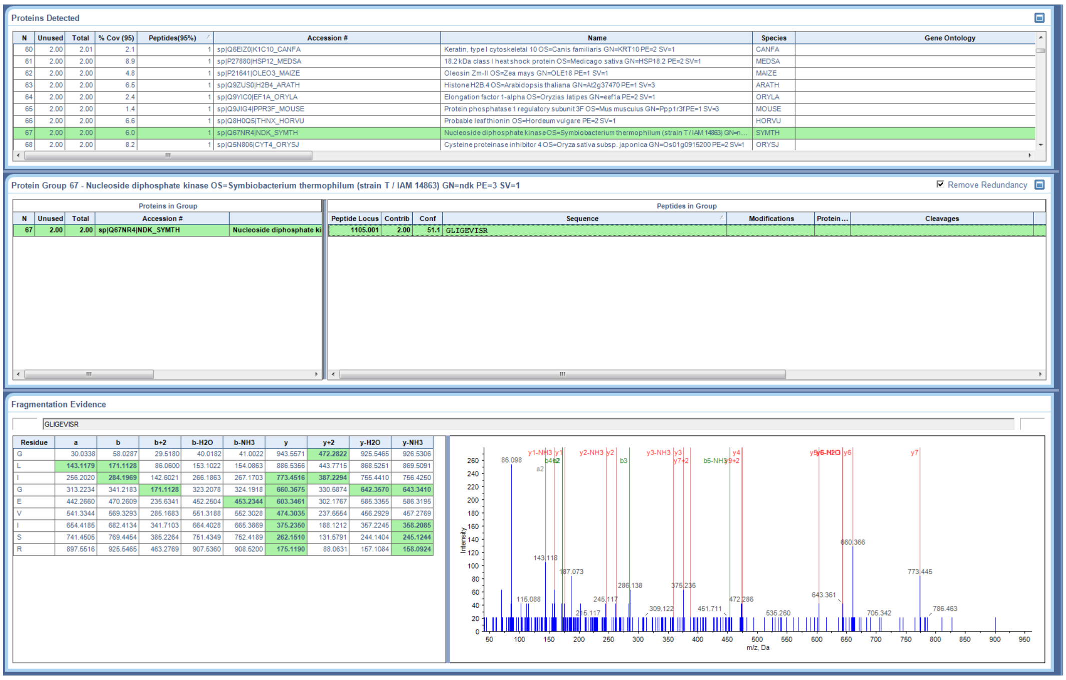


**Supplementary Figure S8. MS/MS of GLIGEVISR identifying NDK_SYMTH.**

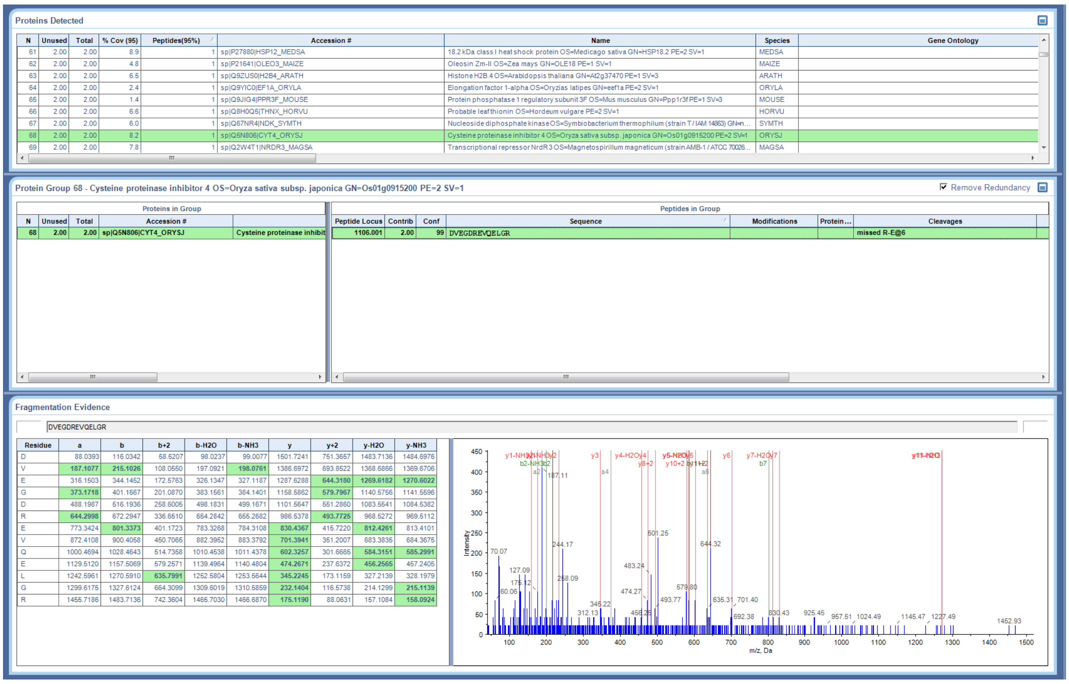


**Supplementary Figure S9. MS/MS of DVEGDREVQELGR identifying CYT4_ORYSJ.**

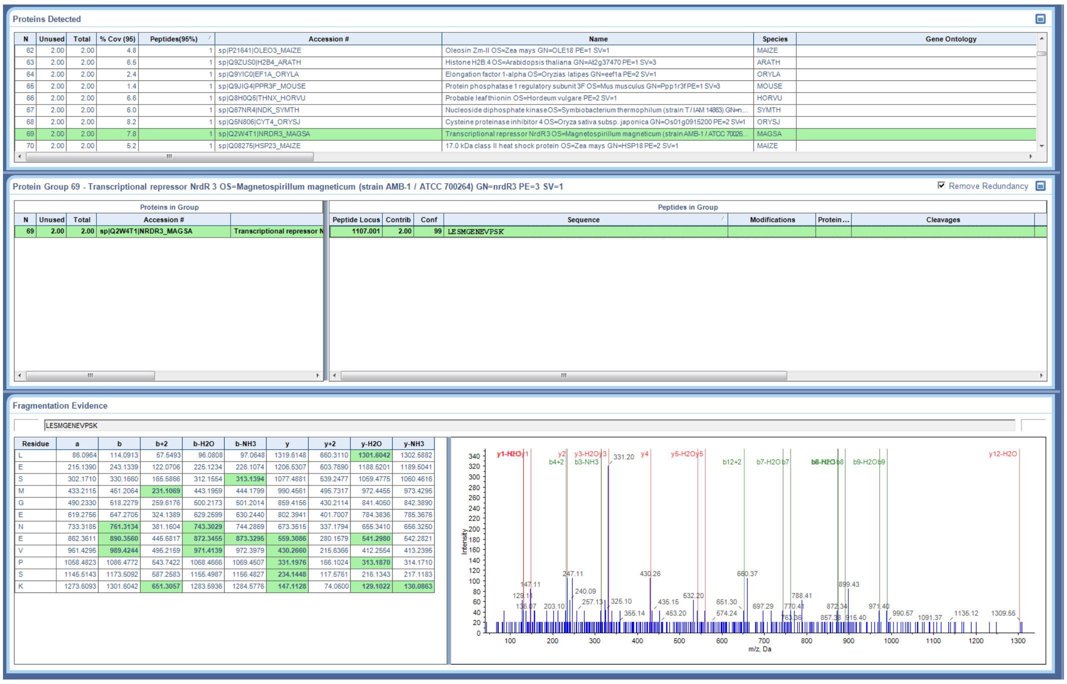


**Supplementary Figure S10. MS/MS of LESMGENEVPSK identifying NRDR3_MAGSA.**

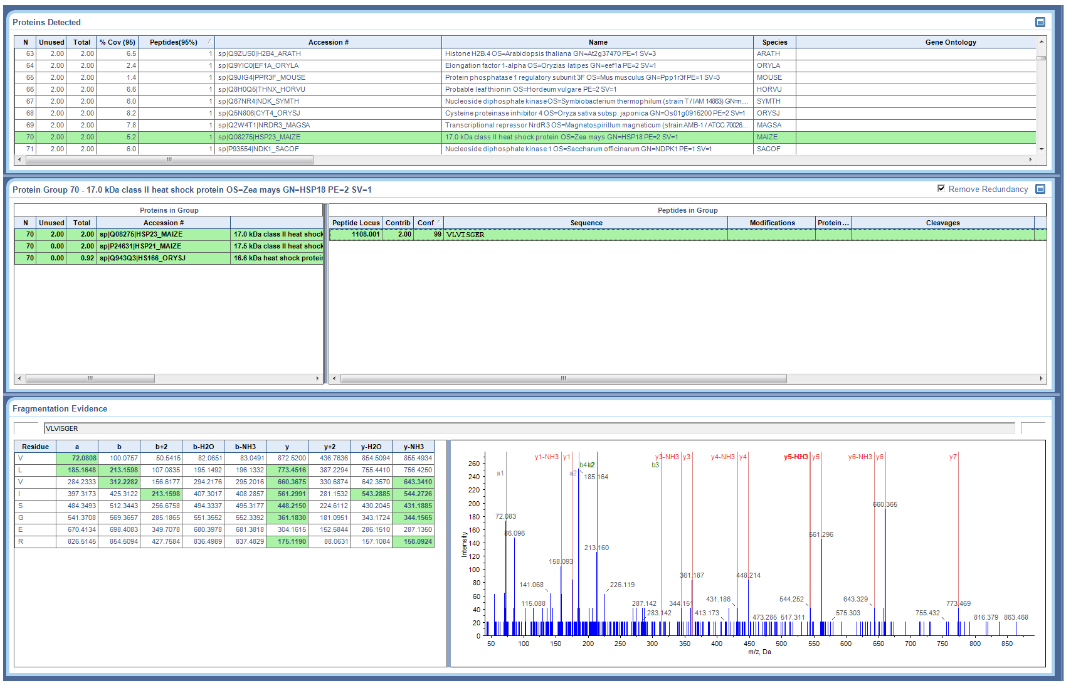


**Supplementary Figure S11. MS/MS of VLVISGER identifying HSP23_MAIZE.**

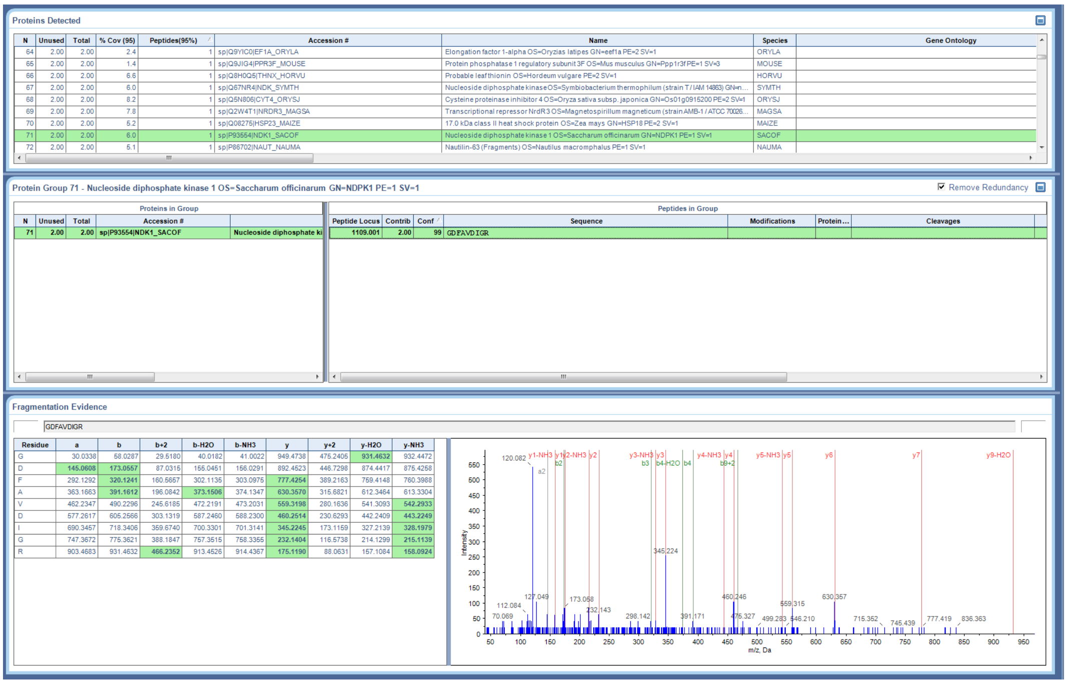


**Supplementary Figure S12. MS/MS of GDFAVDIGR identifying NDK1_SACOF.**

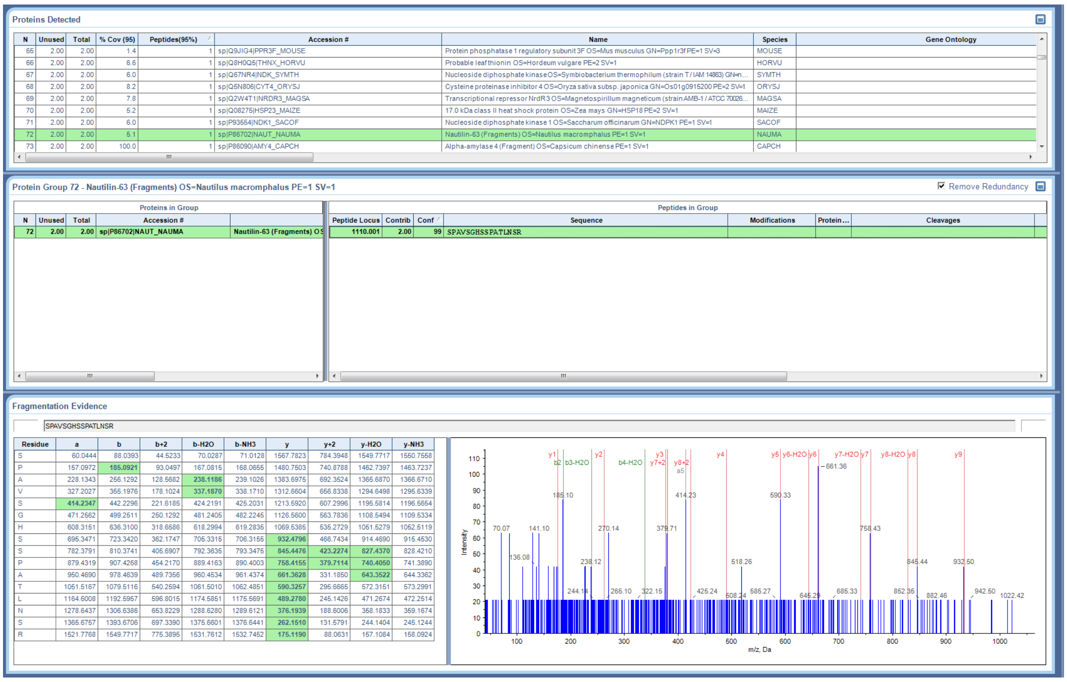


**Supplementary Figure S13. MS/MS of SPAVSGHSSPATLNSR identifying NAUT_NAUMA.**

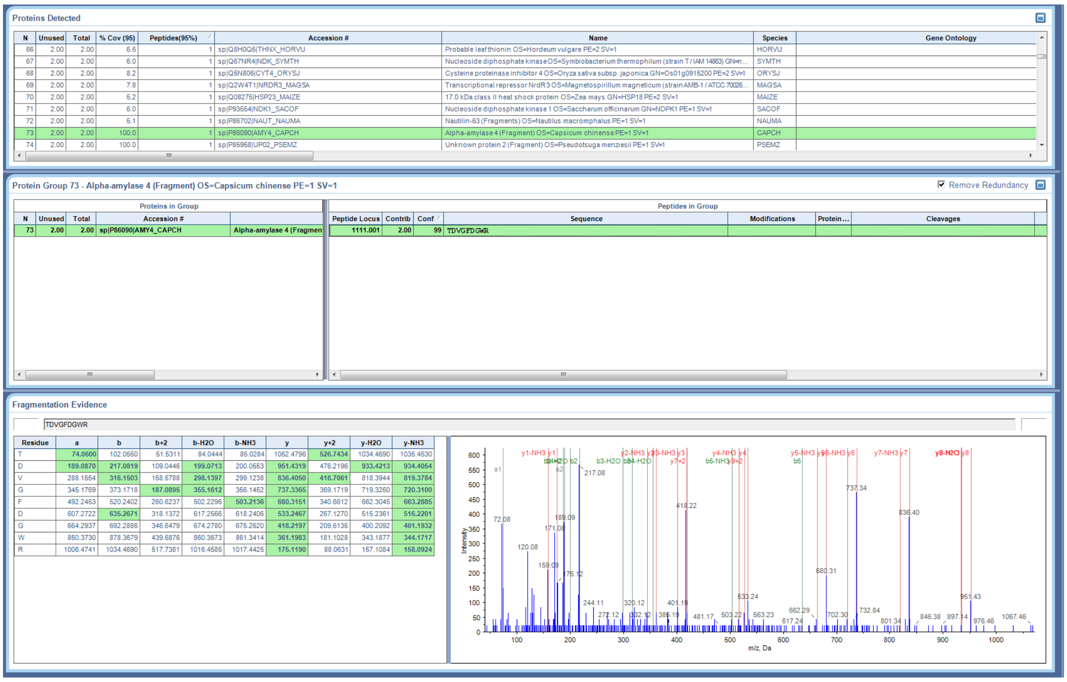


**Supplementary Figure S14. MS/MS of TDVGFDGWR identifying AMY4_CAPCH.**

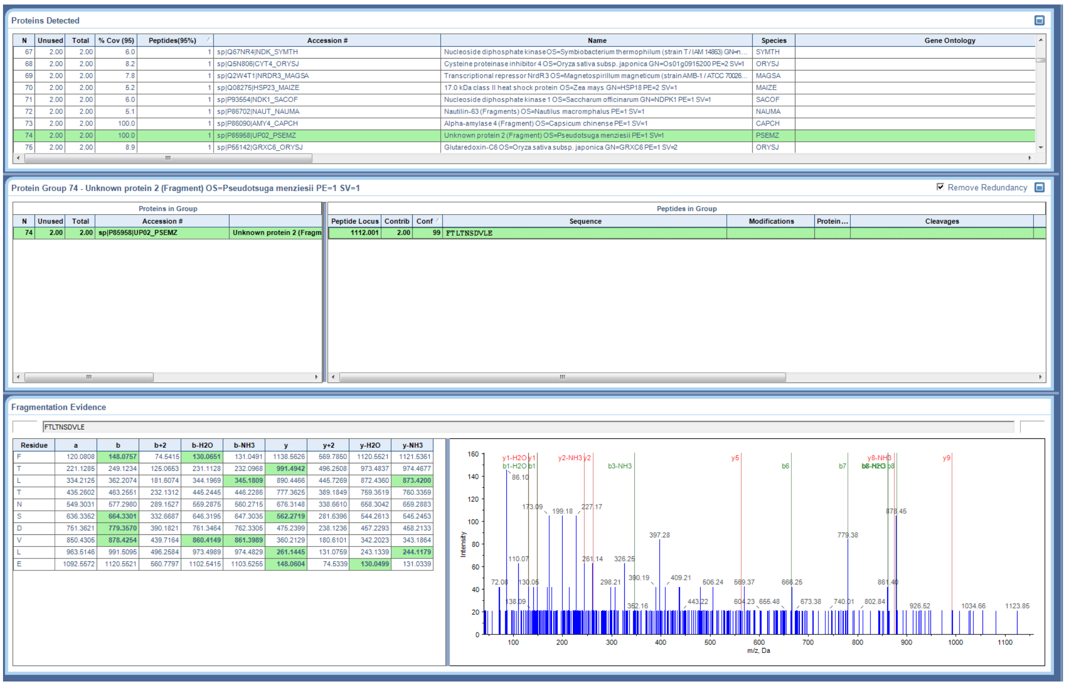


**Supplementary Figure S15. MS/MS of FTLTNSDVLE identifying UP02_PSEMZ.**

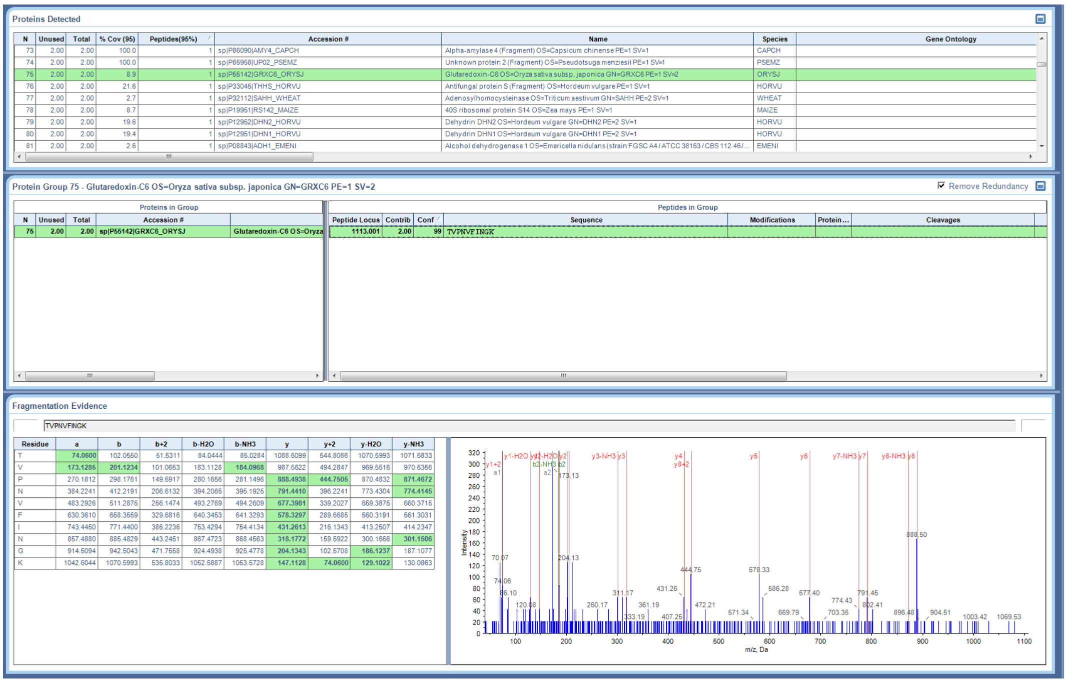


Supplementary Figure S16. MS/MS of TVPNVFINGK identifying GRXC9_ORYSJ.

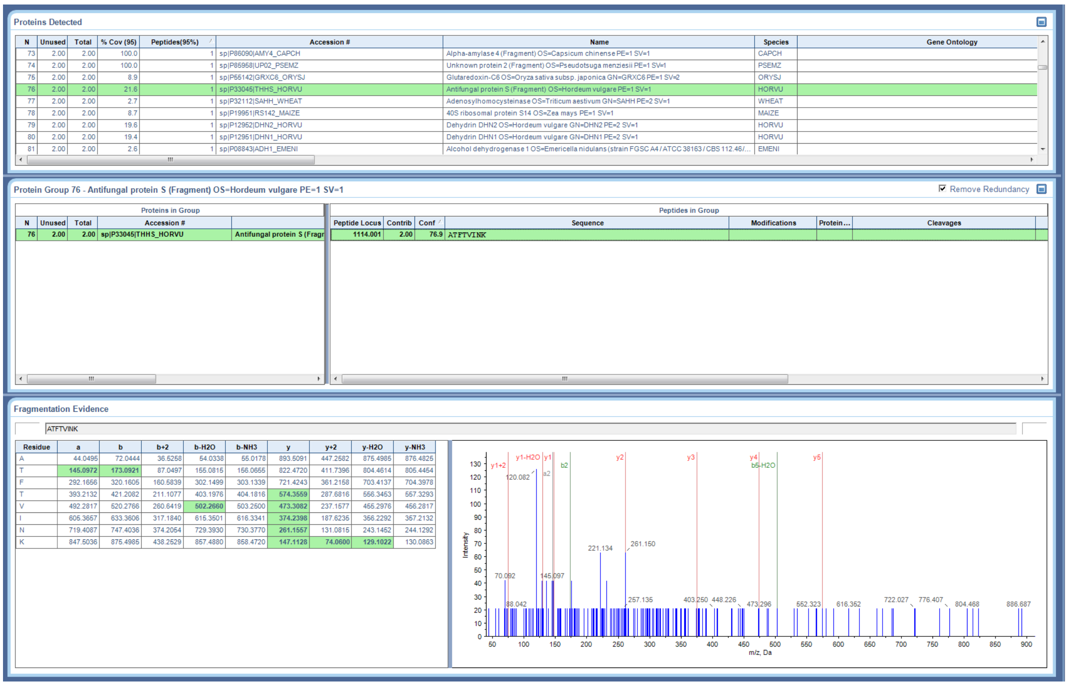


Supplementary Figure S17. MS/MS of ATFTVINK identifying THHS_HORVU.

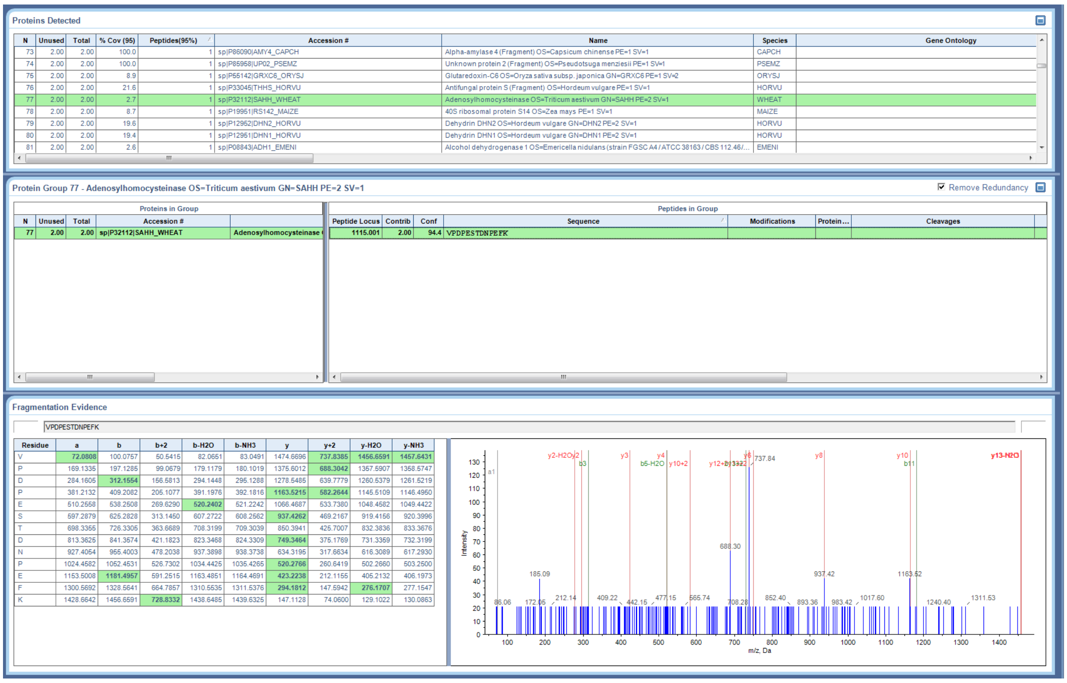


Supplementary Figure S18. MS/MS of VPDPESTDNPEFK identifying SAHH_WHEAT.

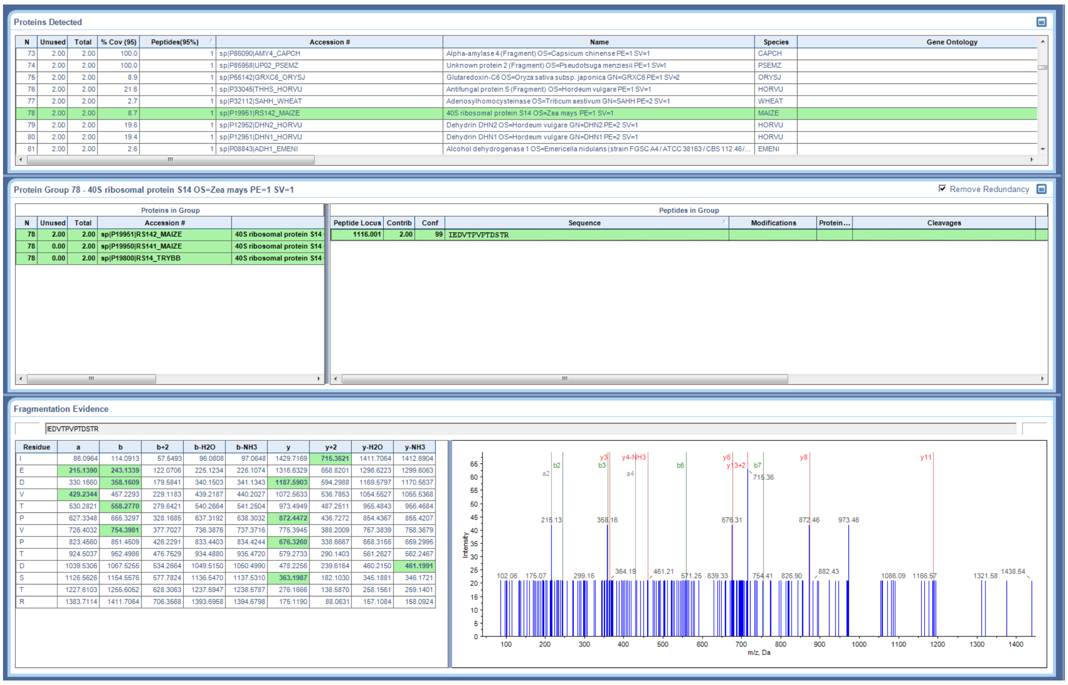


Supplementary Figure S19. MS/MS of IEDVTPVPTDSTR identifying RSI42_MAIZE.

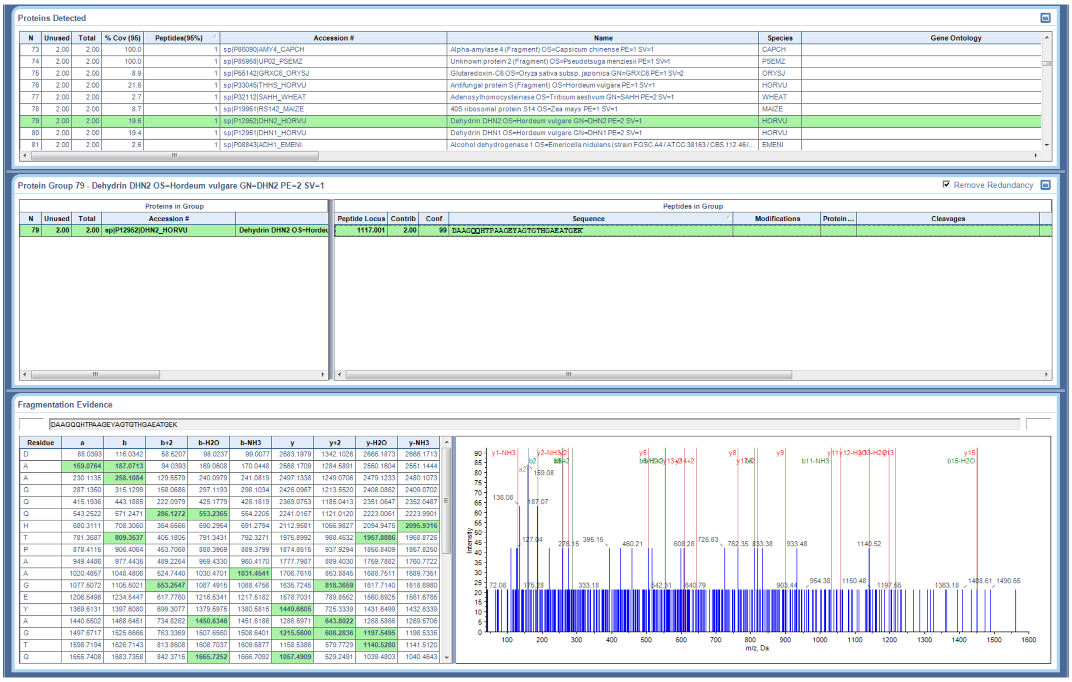


**Supplementary Figure S20. MS/MS of DAAGQQHTPAAGEYAGTGTHGAEATGEK identifying DHN2_HORVU.**

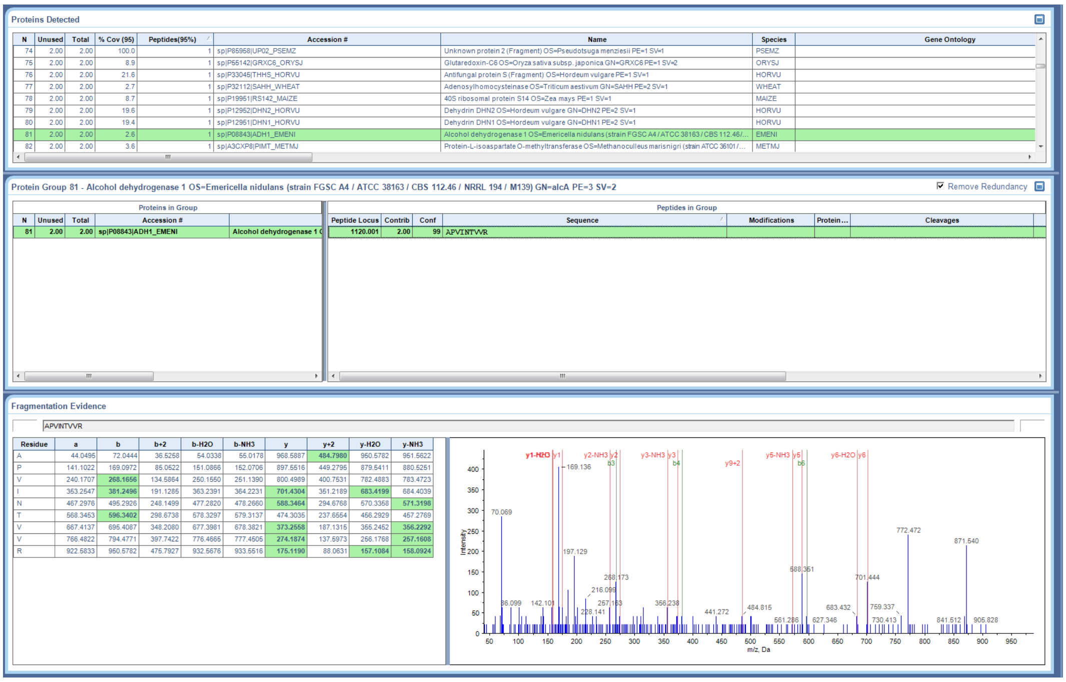


**Supplementary Figure S21. MS/MS of APVINTVVR identifying ADH1_EMENI.**

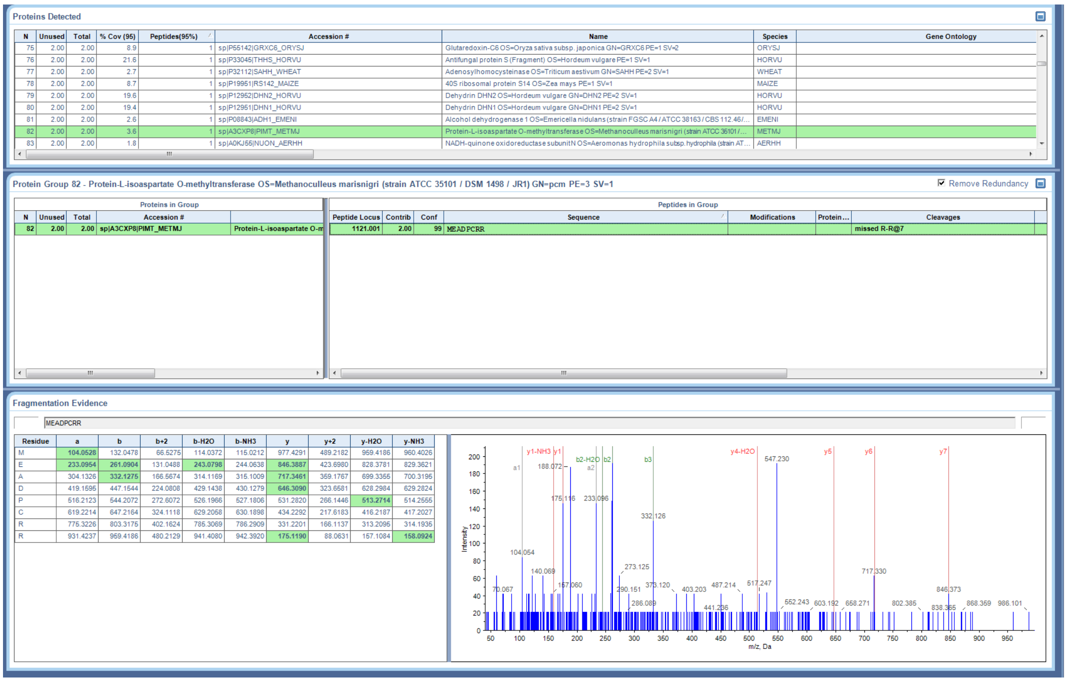


**Supplementary Figure S22. MS/MS of MEADPCRR identifying PIMT_METMJ.**

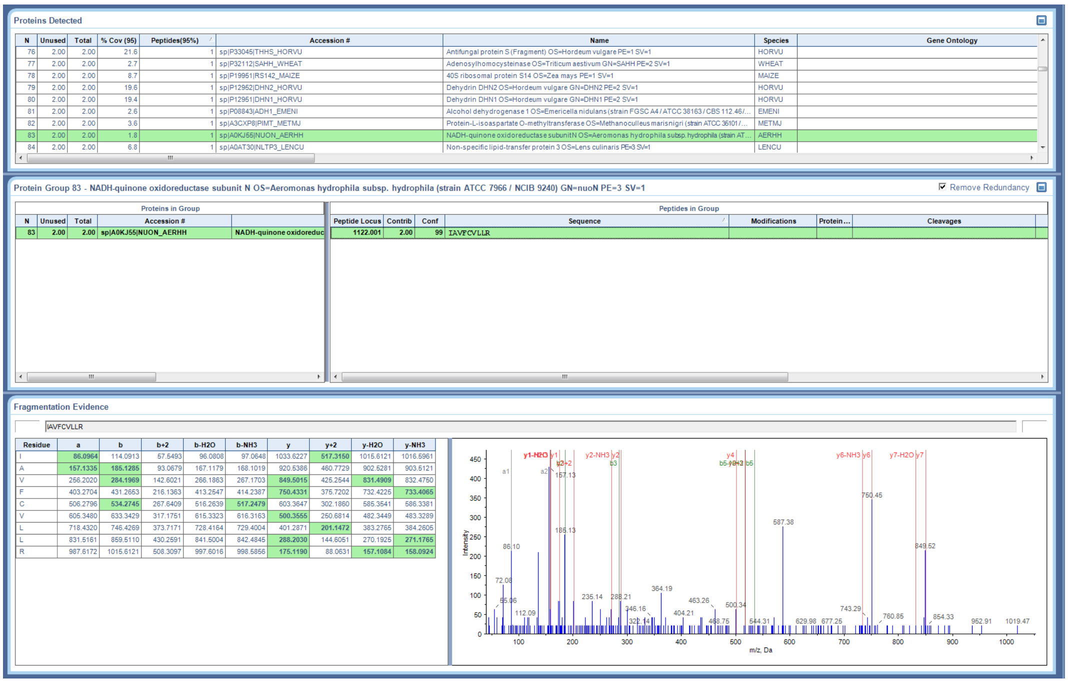


**Supplementary Figure S23. MS/MS of IAVFCVLLR identifying NIJON_AERHH.**

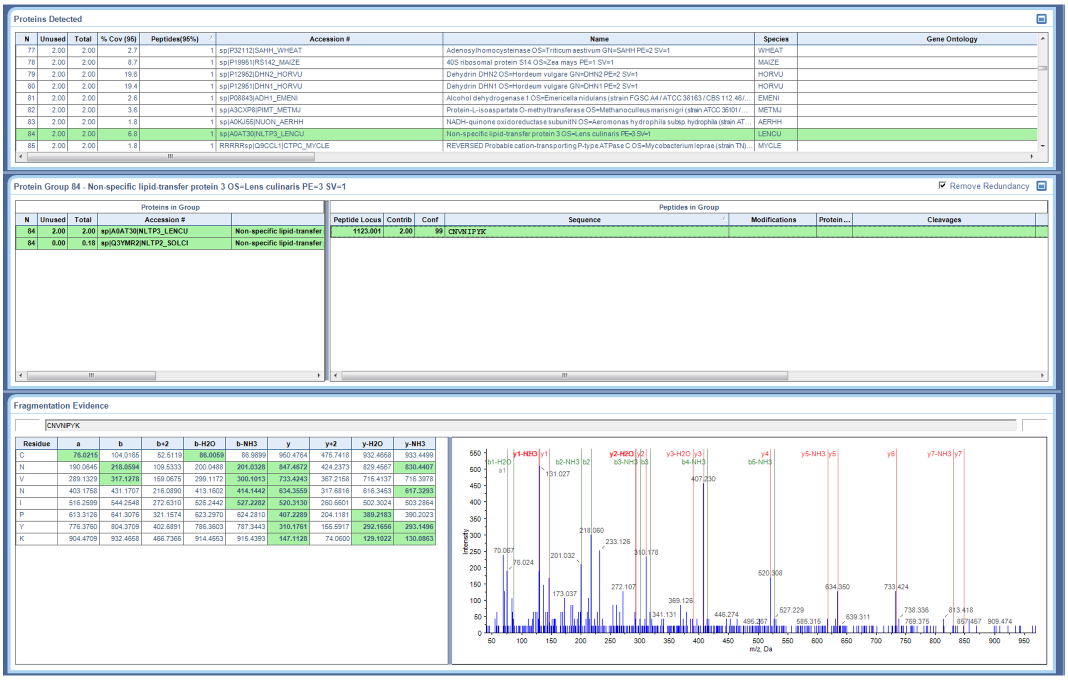


**Supplementary Figure S24. MS/MS of CNVNIPYK identifying NLTP3_LENCU.**

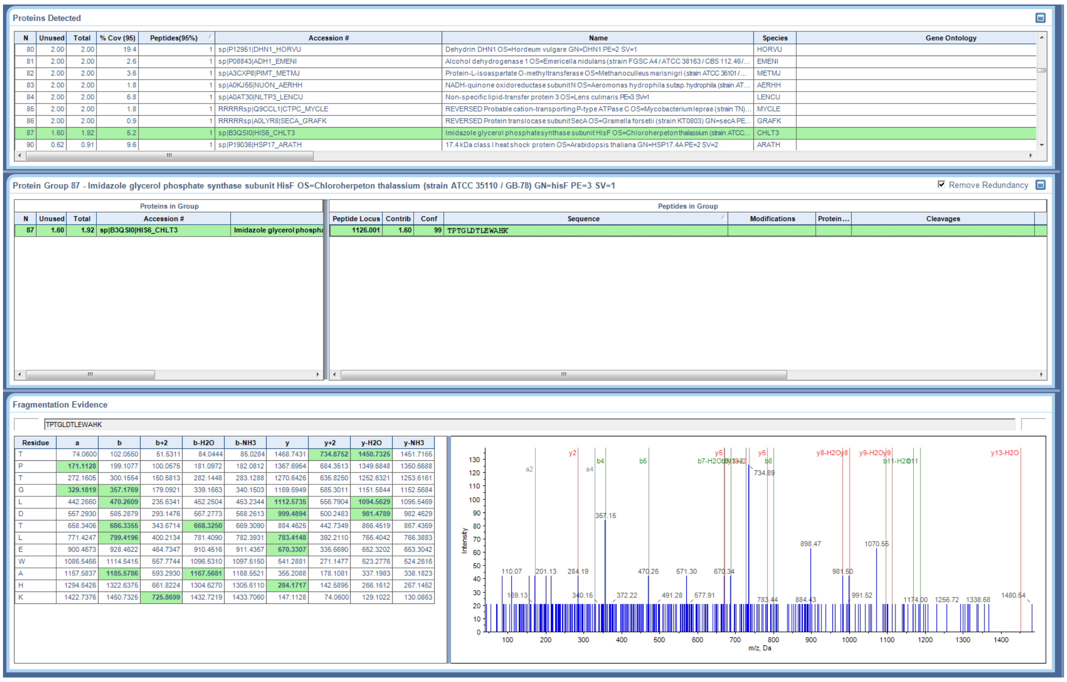


**Supplementary Figure S25. MS/MS of TPTGLDTLEWAHK identifying HIS6_CHLT3.**

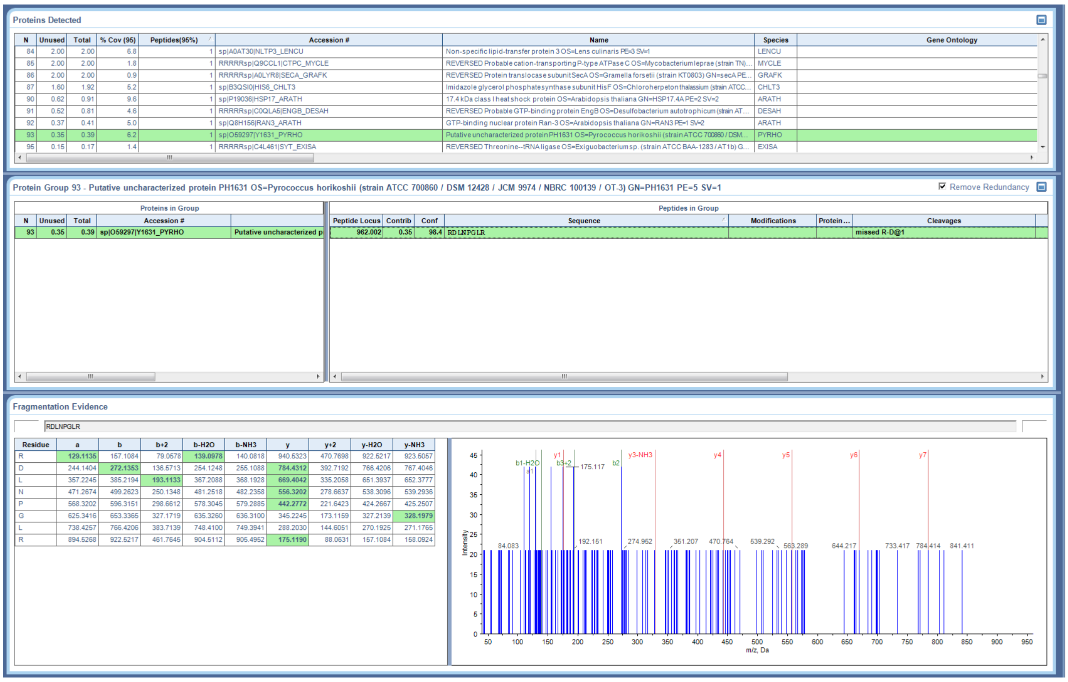


**Supplementary Figure S26. MS/MS of RDLNPGLR identifying Y1631_PYRHO.**

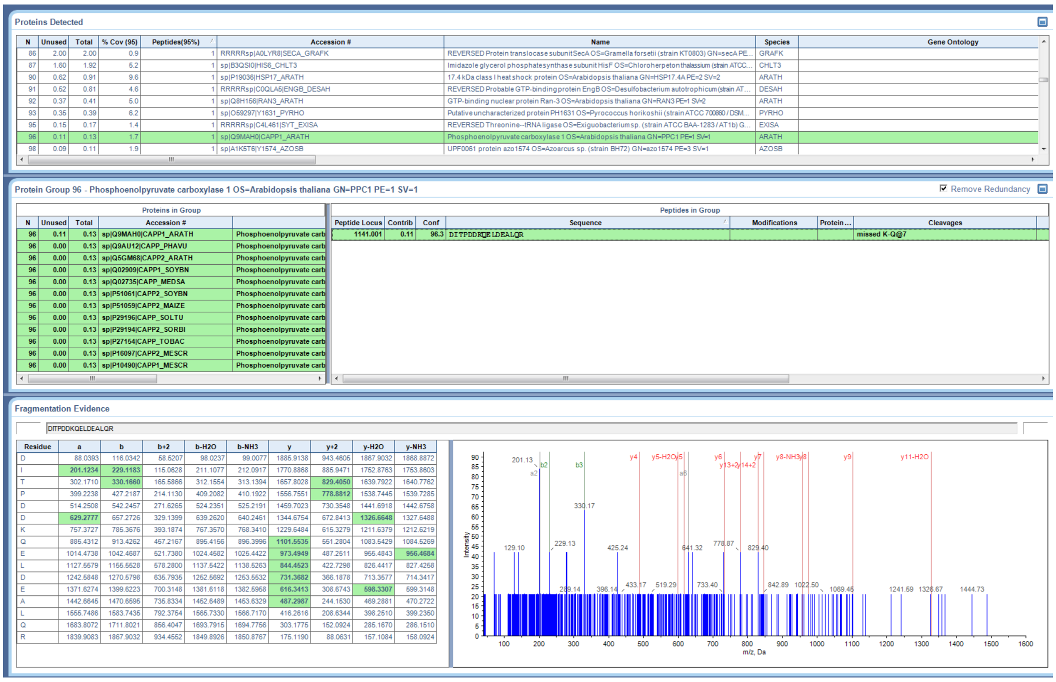


**Supplementary Figure S27. MS/MS of DITPDDKQELDEALQR identifying CAPP1_ARATH.**

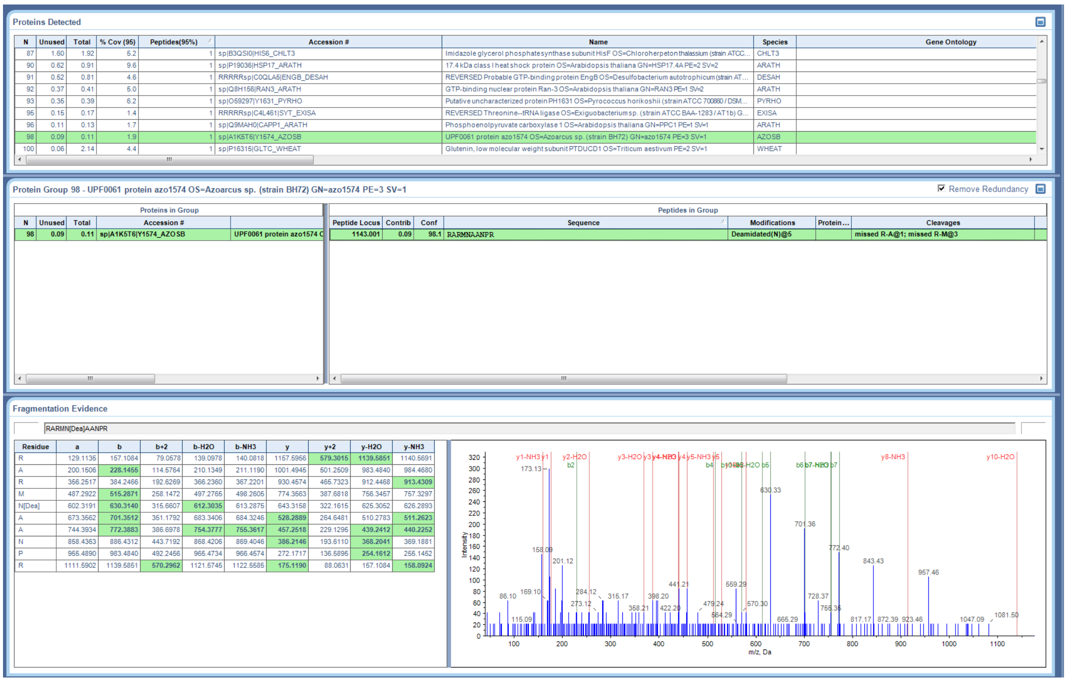


**Supplementary Figure S28. MS/MS of RARMN[Dea]AANPR identifying Y1574_AZOSB.**

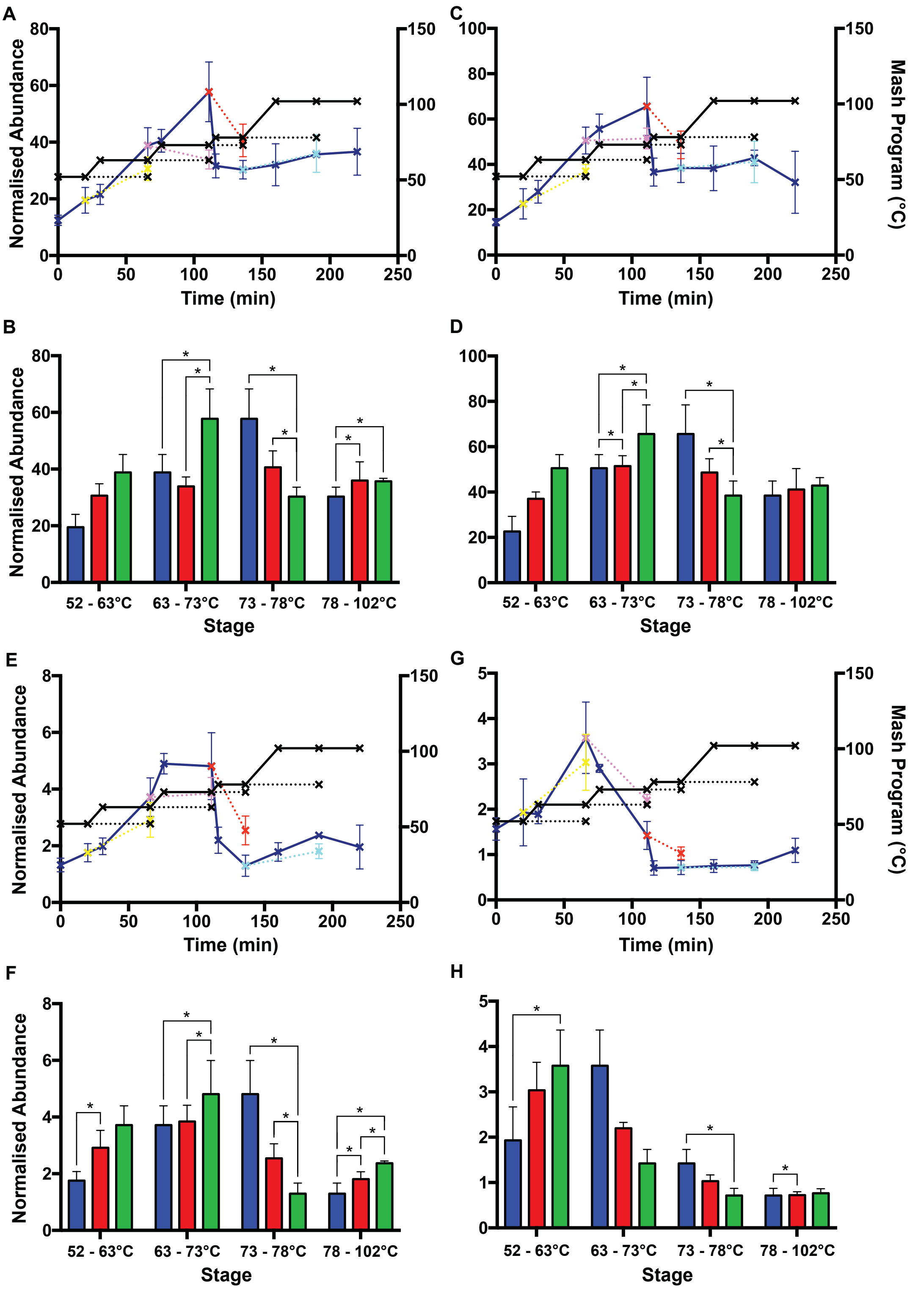


**Supplementary Figure S29. Changes in protein abundance dependent on time or temperature.** Normalized abundance of **(A)** α-amylase/trypsin inhibitor CMb (IAAB), **(C)** α-amylase/trypsin inhibitor CMd (IAAD), **(E)** barwin (BARW), and **(G)** α-amylase type B isozyme (AMY2). Standard mash program, solid line, dark blue; rest stage temperature extensions, dotted lines, yellow (52°C), purple (63°C), red (73°C), and *cyan* (78°C). Mash temperature profile is shown on the right Y-axis. Mash program is shown in black solid and dotted lines; crosses indicate when samples were taken. Normalized abundance of **(B)** IAAB, **(D)** IAAD, **(F)** BARW, and **(H)** AMY2 between sequential stages. First stage (blue), time extension of first stage (red), and next temperature stage (green). *, $P < 0.05$. Values show mean, n=3. Error bars show SEM.

**Supplementary Table 1. Temperature stages used in the mash.**

| Stage | Temperature (°C) | Length of stage (min) |
|---|---|---|
| Protein rest | 52°C | 20 min |
| Maltose rest | 63°C | 35 min |
| Sugar rest 1 | 73°C | 35 min |
| Sugar rest 2 | 78°C | 20 min |
| Boil | 102°C | 60 min |

**Supplementary Table 2. GO term enrichment for the four protein clusters identified by correlation profiling of protein abundance throughout the mash.** Ontology indicates the type of GO term: biological process (BP), molecular function (MF), or chemical component (CC). All GO terms shown were considered significant ($p < 0.05$). GO term enrichment was performed with GOstats.

| GO ID | Ontology | P value | Term | Gene IDs |
|---|---|---|---|---|
| **Cluster 1** | | | | |
| GO:0008289 | MF | 0.003 | lipid binding | NLTP1_HORVU;NLTP2_HORVU;NLTP3_LENCU;NLTP3_WHEAT |
| GO:0010876 | BP | 0.031 | lipid localization | NLTP1_HORVU;NLTP3_LENCU;NLTP3_WHEAT |
| GO:0006869 | BP | 0.031 | lipid transport | NLTP1_HORVU;NLTP3_LENCU;NLTP3_WHEAT |
| GO:0006810 | BP | 0.036 | transport | NLTP1_HORVU;NLTP2_HORVU;NLTP3_LENCU;NLTP3_WHEAT |
| GO:0051234 | BP | 0.036 | establishment of localization | NLTP1_HORVU;NLTP2_HORVU;NLTP3_LENCU;NLTP3_WHEAT |
| GO:0051179 | BP | 0.036 | localization | NLTP1_HORVU;NLTP2_HORVU;NLTP3_LENCU;NLTP3_WHEAT |
| GO:0044092 | BP | 0.046 | negative regulation of molecular function | IAA1_HORVU;IAAE_HORVU |
| GO:0043086 | BP | 0.046 | negative regulation of catalytic activity | IAA1_HORVU;IAAE_HORVU |
| **Cluster 2** | | | | |
| GO:0030234 | MF | 0.001 | enzyme regulator activity | BSZ7_HORVU;CYT4_ORYSJ;HINB2_HORVU;IAA2_HORVU;IAAA_HORVU;IAAB_HORVU;IAAD_HORVU;IAA_HORVU;IAC16_WHEAT;ICIA_HORVU;ICIB_HORVU |
| GO:0098772 | MF | 0.001 | molecular function regulator | BSZ7_HORVU;CYT4_ORYSJ;HINB2_HORVU;IAA2_HORVU;IAAA_HORVU;IAAB_HORVU;IAAD_HORVU;IAA_HORVU;IAC16_WHEAT;ICIA_HORVU;ICIB_HORVU |
| GO:0004866 | MF | 0.001 | endopeptidase inhibitor activity | BSZ7_HORVU;CYT4_ORYSJ;HINB2_HORVU;IAA2_HORVU;IAAA_HORVU;IAAB_HORVU;IAAD_HORVU;IAA_HORVU;IAC16_WHEAT;ICIA_HORVU;ICIB_HORVU |
| GO:0004857 | MF | 0.001 | enzyme inhibitor activity | BSZ7_HORVU;CYT4_ORYSJ;HINB2_HORVU;IAA2_HORVU;IAAA_HORVU;IAAB_HORVU;IAAD_HORVU;IAA_HORVU;IAC16_WHEAT;ICIA_HORVU;ICIB_HORVU |
| GO:0030414 | MF | 0.001 | peptidase inhibitor activity | BSZ7_HORVU;CYT4_ORYSJ;HINB2_HORVU;IAA2_HORVU;IAAA_HORVU;IAAB_HORVU;IAAD_HORVU;IAA_HORVU;IAC16_WHEAT;ICIA_HORVU;ICIB_HORVU |
| GO:0061134 | MF | 0.001 | peptidase regulator activity | BSZ7_HORVU;CYT4_ORYSJ;HINB2_HORVU;IAA2_HORVU;IAAA_HORVU;IAAB_HORVU;IAAD_HORVU;IAA_HORVU;IAC16_WHEAT;ICIA_HORVU;ICIB_HORVU |
| GO:0061135 | MF | 0.001 | endopeptidase regulator activity | BSZ7_HORVU;CYT4_ORYSJ;HINB2_HORVU;IAA2_HORVU;IAAA_HORVU;IAAB_HORVU;IAAD_HORVU;IAA_HORVU;IAC16_WHEAT;ICIA_HORVU;ICIB_HORVU |
| GO:0004867 | MF | 0.003 | serine-type endopeptidase inhibitor activity | BSZ7_HORVU;HINB2_HORVU;IAA2_HORVU;IAAA_HORVU;IAAB_HORVU;IAAD_HORVU;IAA_HORVU;IAC16_WHEAT;ICIA_HORVU;ICIB_HORVU |
| GO:0015066 | MF | 0.013 | alpha-amylase inhibitor activity | IAA2_HORVU;IAAA_HORVU;IAAB_HORVU;IAAD_HORVU;IAA_HORVU;IAC16_WHEAT |
| GO:0006139 | BP | 0.049 | nucleobase-containing compound metabolic process | ALF2_CAEEL;NDK1_SACOF;NRDR3_MAGSA;TPIS_HORVU |
| **Cluster 3** | | | | |
| GO:0016798 | MF | 0.000 | hydrolase activity, acting on glycosyl bonds | AMY2_HORVU;AMY4_CAPCH;AMY6_HORVU;AMYB_HORVS;AMYB_HORVU;BGL26_ORYSJ;CHI1_HORVU;CHI2_HORVU |
| GO:0004553 | MF | 0.000 | hydrolase activity, hydrolyzing O-glycosyl compounds | AMY2_HORVU;AMY4_CAPCH;AMY6_HORVU;AMYB_HORVS;AMYB_HORVU;BGL26_ORYSJ;CHI1_HORVU;CHI2_HORVU |
| GO:0016787 | MF | 0.000 | hydrolase activity | AMY2_HORVU;AMY4_CAPCH;AMY6_HORVU;AMYB_HORVS;AMYB_HORVU;BGL26_ORYSJ;CHI1_HORVU;CHI2_HORVU;CTPC_MYCLE |
| GO:0003824 | MF | 0.000 | catalytic activity | AMY2_HORVU;AMY4_CAPCH;AMY6_HORVU;AMYB_HORVS;AMYB_HORVU;BGL26_ORYSJ;CHI1_HORVU;CHI2_HORVU;CTPC_MYCLE;GRDH_ORYSJ;LGUL_ORYSJ;NUON_AERHH |
| GO:0016160 | MF | 0.001 | amylase activity | AMY2_HORVU;AMY4_CAPCH;AMY6_HORVU;AMYB_HORVS;AMYB_HORVU |

| | | | | |
|---|---|---|---|---|
| GO:0004556 | MF | 0.019 | alpha-amylase activity | AMY2_HORVU;AMY4_CAPCH;AMY6_HORVU |
| GO:0004568 | MF | 0.032 | chitinase activity | CHI1_HORVU;CHI2_HORVU |
| GO:0016161 | MF | 0.032 | beta-amylase activity | AMYB_HORVS;AMYB_HORVU |
| GO:0005975 | BP | 0.000 | carbohydrate metabolic process | AMY2_HORVU;AMY4_CAPCH;AMY6_HORVU;AMYB_HORVS;AMYB_HORVU;BGL26_ORYSJ;CHI1_HORVU;CHI2_HORVU |
| GO:0071704 | BP | 0.000 | organic substance metabolic process | AMY2_HORVU;AMY4_CAPCH;AMY6_HORVU;AMYB_HORVS;AMYB_HORVU;BGL26_ORYSJ;CHI1_HORVU;CHI2_HORVU;NUON_AERHH |
| GO:0044238 | BP | 0.000 | primary metabolic process | AMY2_HORVU;AMY4_CAPCH;AMY6_HORVU;AMYB_HORVS;AMYB_HORVU;BGL26_ORYSJ;CHI1_HORVU;CHI2_HORVU;NUON_AERHH |
| GO:0009057 | BP | 0.000 | macromolecule catabolic process | AMYB_HORVS;AMYB_HORVU;CHI1_HORVU;CHI2_HORVU |
| GO:0000272 | BP | 0.000 | polysaccharide catabolic process | AMYB_HORVS;AMYB_HORVU;CHI1_HORVU;CHI2_HORVU |
| GO:0008152 | BP | 0.001 | metabolic process | AMY2_HORVU;AMY4_CAPCH;AMY6_HORVU;AMYB_HORVS;AMYB_HORVU;BGL26_ORYSJ;CHI1_HORVU;CHI2_HORVU;NUON_AERHH |
| GO:0005976 | BP | 0.002 | polysaccharide metabolic process | AMYB_HORVS;AMYB_HORVU;CHI1_HORVU;CHI2_HORVU |
| GO:0016052 | BP | 0.020 | carbohydrate catabolic process | AMYB_HORVS;AMYB_HORVU;CHI1_HORVU;CHI2_HORVU |
| GO:0006022 | BP | 0.025 | aminoglycan metabolic process | CHI1_HORVU;CHI2_HORVU |
| GO:0006026 | BP | 0.025 | aminoglycan catabolic process | CHI1_HORVU;CHI2_HORVU |
| GO:0006040 | BP | 0.025 | amino sugar metabolic process | CHI1_HORVU;CHI2_HORVU |
| GO:0006030 | BP | 0.025 | chitin metabolic process | CHI1_HORVU;CHI2_HORVU |
| GO:0006032 | BP | 0.025 | chitin catabolic process | CHI1_HORVU;CHI2_HORVU |
| GO:0016998 | BP | 0.025 | cell wall macromolecule catabolic process | CHI1_HORVU;CHI2_HORVU |
| GO:0071554 | BP | 0.025 | cell wall organization or biogenesis | CHI1_HORVU;CHI2_HORVU |
| GO:0046348 | BP | 0.025 | amino sugar catabolic process | CHI1_HORVU;CHI2_HORVU |
| GO:0044036 | BP | 0.025 | cell wall macromolecule metabolic process | CHI1_HORVU;CHI2_HORVU |
| GO:1901136 | BP | 0.025 | carbohydrate derivative catabolic process | CHI1_HORVU;CHI2_HORVU |
| GO:1901071 | BP | 0.025 | glucosamine-containing compound metabolic process | CHI1_HORVU;CHI2_HORVU |
| GO:1901072 | BP | 0.025 | glucosamine-containing compound catabolic process | CHI1_HORVU;CHI2_HORVU |
| GO:1901575 | BP | 0.033 | organic substance catabolic process | AMYB_HORVS;AMYB_HORVU;CHI1_HORVU;CHI2_HORVU |
| GO:0009056 | BP | 0.033 | catabolic process | AMYB_HORVS;AMYB_HORVU;CHI1_HORVU;CHI2_HORVU |
| GO:0044425 | CC | 0.004 | membrane part | CTPC_MYCLE;NUON_AERHH |
| GO:0031224 | CC | 0.004 | intrinsic component of membrane | CTPC_MYCLE;NUON_AERHH |
| GO:0016021 | CC | 0.004 | integral component of membrane | CTPC_MYCLE;NUON_AERHH |
| GO:0071944 | CC | 0.007 | cell periphery | CTPC_MYCLE;NUON_AERHH |
| GO:0005886 | CC | 0.007 | plasma membrane | CTPC_MYCLE;NUON_AERHH |
| GO:0016020 | CC | 0.033 | membrane | CTPC_MYCLE;NUON_AERHH |
| **Cluster 4** | | | | |
| GO:0003824 | MF | 0.032 | catalytic activity | AMY1_HORVU;CYSP2_HORVU;G3PC1_HORVU;HIS6_CHLT3;PDI_WHEAT;PIMT_METMJ;REHY_HORVU;SODC2_ORYSJ |
| GO:0044267 | BP | 0.023 | cellular protein metabolic process | PIMT_METMJ;RS142_MAIZE;RUB1_ARATH |
| GO:0005737 | CC | 0.048 | cytoplasm | G3PC1_HORVU;HIS6_CHLT3;PDI_WHEAT;PIMT_METMJ;RS142_MAIZE;RUB1_ARATH;SODC2_ORYSJ |

[1] Biological Process
[2] Molecular Function
[3] Cellular Compartment

**Supplementary Table 3. ProteinPilot identification summary.**

**Supplementary Table 4. PeakView SWATH summary.**

**Supplementary Table 5. Multiple comparisons of mash steps via Tukey contrasts following one-way ANOVA on each protein over steps of mash.** $P < 0.05$ indicates significance between two steps.

**Supplementary Table 6. Multiple Comparisons of Means via Tukey Contrasts following one-way ANOVA on each analysed peptide over steps of mash.** $P < 0.05$ indicates significance between two steps.

**Supplementary Table 7. Temperature stages used in the micro-mash.**

| Stage | Temperature (°C) | Length of stage (min) |
|---|---|---|
| Protein rest | 52°C | 20 min |
| Protein rest extension | | 35 min (length of maltose rest) |
| Maltose rest | 63°C | 35 min |
| Maltose rest extension | | 35 min (length of sugar rest 1) |
| Sugar rest 1 | 73°C | 35 min |
| Sugar rest 1 extension | | 20 min (length of sugar rest 2) |
| Sugar rest 2 | 78°C | 20 min |
| Sugar rest 2 extension | | 30 min (length of half the boil) |
| Boil | 102°C | 60 min |

**Supplementary Material 1. PeakView SWATH file.**
